# Neural mechanisms of handedness for precision drawing: hand-dependent engagement of cortical networks for bimanual control

**DOI:** 10.1101/2025.11.18.689091

**Authors:** Namarta Kapil, Taewon Kim, Samah Gassass, Ruiwen Zhou, Alexandre R. Carter, Ian G. Dobbins, Lei Liu, Mark P. McAvoy, Muriah D. Wheelock, Yong Wang, David M. Brogan, Christopher J. Dy, Susan E. Mackinnon, Benjamin A. Philip

**Author notes:** Corresponding Author: Benjamin Philip. Co-first authors (Namarta Kapil, Taewon Kim).

## Abstract

Neural mechanisms underlying handedness remain poorly understood. We used functional magnetic resonance imaging (fMRI) to study performance of a visually guided drawing task with each hand. We hypothesized that the left superior parietal lobule supports drawing with either hand, and individuals with chronic peripheral nerve injury (PNI) to the dominant hand use the same mechanism as healthy adults. Thirty-three right-handed adults (23 healthy, 10 patients) underwent fMRI while performing a precision drawing task, alternating between the right hand (RH) and left hand (LH). BOLD magnitude and functional connectivity (FC) modulation via generalized psychophysiological interaction were analyzed in 12 a priori regions of interest, followed by additional exploratory areas identified via whole-brain magnitude analysis. During LH drawing (compared to RH drawing), contralateral primary motor cortex showed lower BOLD magnitude but greater FC with two networks: First, a left motor-premotor network with increased FC and equal-or-greater magnitude during LH drawing. Second, a right parietal network characterized by increased FC during LH drawing but increased magnitude during RH drawing. Exploratory whole-brain analyses supported and extended these motor and parietal networks, and additionally suggested that RH drawing may involve greater magnitude and FC within a bilateral parieto-premotor network centered on right paracingulate cortex. No significant group-related effects were observed in BOLD magnitude or FC. These results describe the first proposed mechanisms for LH precision drawing, both of which depend on differential engagement of bimanual control networks: a left hemisphere motor-premotor network for precision motor control, which engages intrahemispherically (directly) during RH drawing and interhemispherically (indirectly) during LH drawing; and a right hemisphere parietal network with greater distributed coordination during LH drawing. Although these mechanisms’ preservation after PNI requires confirmation in a larger sample, their differential involvement in LH precision drawing supports their potential relevance as neuromodulatory targets to enhance LH performance after RH impairment.

## 1. INTRODUCTION

Human handedness is a multifactorial construct (Dexheimer et al., 2024), one aspect of which is left-right asymmetries in upper limb performance (“performance asymmetries” for brevity). Little is known about the neural mechanisms underlying these asymmetries, even though they are routinely expressed during our everyday actions. According to the complementary dominance theory (Kitchen et al., 2025; Shabbott,B.A., & Sainburg, R.L., 2008), these asymmetries arise from lateralized cortical specializations for different aspects of movement: in right-handed individuals, the left hemisphere contains specializations for precision control of trajectories and predictive dynamics, while the right hemisphere contains specializations for stabilization and impedance control. Each of these lateralized specializations supports bilateral action, but is primarily connected to the contralateral hand, leading to e.g. a right-hand (RH) advantage for smooth trajectories. However, these neural specializations cannot yet be assessed or targeted to support rehabilitation because the specific neural mechanisms of performance asymmetries remain unknown.

Our current understanding of the neural mechanisms underlying performance asymmetries is based on lesion and neuroimaging studies, each of which offer different limitations. Lesion studies have shown that the left hemisphere contains a specialization that supports precision trajectory control with both hands, and is expressed across motor tasks with varying demands, including two-dimensional reaching tasks with complex demands (Schaefer et al., 2012), and simpler tasks such as ballistic two-dimensional reaching (Mani et al., 2013; Schaefer et al., 2009) and finger-tapping sequences (Haaland et al., 2004).

However, lesion studies cannot provide anatomical precision due to lesion heterogeneity, nor can they distinguish between stroke-related deficits and post-stroke adaptation. Neuroimaging allows precise spatial measurement, but is often restricted to simpler movements: such studies have identified a role for left M1 in bilateral finger tapping (Hayashi et al., 2008; Tzourio-Mazoyer et al., 2015), increased involvement of ipsilateral M1 under high task demands during two-finger joystick movements (Barany et al., 2020; Buetefisch et al., 2014; Wischnewski et al., 2016), and greater interhemispheric functional connectivity (FC) for the non-preferred hand (NPH) vs. preferred hand (PH) during simple open/close fist movements (Tsurugizawa et al., 2023). Overall, these studies provide important insights but remain limited in their ability to capture the full scope of real-world movement control, as few neuroimaging tasks engage the precision trajectory control that defines the PH’s performance advantage, let alone the mechanisms involved in real-world upper limb actions.

Real-world movements—such as writing, using utensils, or reaching in cluttered environments—involve a dynamic integration of feedforward and feedback mechanisms (Kringelbach et al., 2023; Peterson et al., 2021). Investigating the neural basis of these real-world processes necessitates ecologically valid tasks that capture the complexity and variability of natural behavior rather than artificially constrained experimental conditions (Finn et al., 2022; Finn et al., 2020) that predominate much of the existing lesion and neuroimaging literature. This recognition has prompted a growing shift in cognitive neuroscience toward naturalistic functional MRI (fMRI) designs that incorporate dynamic stimuli such as movies, narratives, and virtual reality (Bottenhorn et al., 2018; Saarimaki, 2021; Zhang et al., 2021). These approaches elicit rich, multimodal brain activity and engage distributed neural networks, including regions that remain unrecruited during simpler tasks, leading to deeper insights into real-world cognition (Hasson et al., 2010). Extending this naturalistic approach to motor control research, however, remains challenging in the fMRI environment due to postural constraints and motion-related artifacts (Reddy et al., 2024; Scarapicchia et al., 2017). Despite these challenges, naturalistic motor tasks have been successfully used to elicit distinct activation patterns via real-time visuomotor feedback (Karimpoor et al., 2018; Yang et al., 2024) and reaching and grasping of 3D objects (Freud et al., 2018). These findings support the need for demanding and ecologically-relevant motor paradigms in fMRI studies to understand the neural basis of lateralized motor control.

The neural mechanisms of handedness hold critical clinical relevance to patients with chronic impairment of the PH. Chronic PH impairment occurs frequently despite surgery and/or rehabilitation, affecting approximately 22% of hemiplegic stroke patients (Stinear et al., 2012) and 22% of patients with upper extremity peripheral nerve injury (PNI) (Dyck et al., 2005; He et al., 2014; Philip et al., 2020). Better outcomes following PH-PNI are associated with improved performance of the NPH (Kim et al., 2024), but strategies to facilitate NPH use during rehabilitation remain poorly understood (Kim et al., 2025; Marcori et al., 2019). Bridging this gap requires a mechanistic understanding of the asymmetric neural mechanisms of naturalistic action. To that end, we developed an fMRI visually guided precision drawing task (Philip & Frey, 2014; Philip et al., 2023) designed to engage participants in an ecologically relevant activity (control of a pen) that requires both feedforward and feedback control during continuous dynamic movements. Critically, pen control skills have a strong PH advantage in most humans (Blank et al., 2000), and therefore are uniquely impaired in patients with chronic impairment of the PH. By having participants perform this task with their PH and NPH, we aim to isolate the mechanisms that support PH-like action in the NPH –mechanisms that could be targeted to enhance rehabilitation outcomes in individuals with chronic PH deficits.

In the current study, we isolated “PH-like action with the NPH” (in right-handed participants, thus “RH-like action with the LH”) via a task fMRI study of participants performing a precision drawing task with each hand (LH and RH, separately). We analyzed two complementary measures of brain function: blood oxygenation level-dependent (BOLD) magnitude, which reflects local task-evoked neural activity; and functional connectivity (FC), which reflects temporally correlated fluctuations in spontaneous neural activity across distributed brain regions (Coito et al., 2019; Fu et al., 2017; Gutierrez-Barragan et al., 2019). We analyzed task-dependent modulation of BOLD magnitude using general linear models (GLM), and task-dependent modulation of FC using generalized psychophysiological interaction (gPPI) (McLaren et al., 2012). Based on previous work (Philip et al., 2021), we hypothesized that precision drawing’s LH > RH difference (hand_LH-RH_) would entail greater ipsilateral BOLD magnitude in human intraparietal area 3 within the superior parietal lobule (SPL_hIP3_). In addition, to establish neural correlates for the processes of adapting to chronic PH impairment after neurological injury, we hypothesized that patients with chronic PNI to the RH would use the same neural mechanisms as healthy adults.

## 2. METHODS

### 2.1 Participants

Thirty-three right-handed adults (24 female; ages 47 ± 18, range 24-82) performed a visually guided precision drawing task (see *fMRI Task* below) in the fMRI scanner. Of these 33 participants, 10 had peripheral nerve injuries (PNI) to their right arm (“patients”; ages 52 ± 17, female 73%), and 23 were healthy controls (ages 46 ± 18, female 74%). The two groups did not differ in age, sex, race/ethnicity, or education (p > 0.4); for full demographics see **Supplementary Table 1**. Patient clinical characteristics are described in **Table 1**. Two additional patients were excluded from the current analysis because they were unable to complete the task with their affected RH, and four additional participants were excluded due to poor task compliance or excess motion as detailed below (see *2.C fMRI Motion Outlier Identification and Screening*). These six participants were therefore not included in the final sample (n = 33). Demographic and performance data were stored and managed via the Research Electronic Data Capture system (Harris et al., 2009). Informed consent was obtained from all participants for being included in the study, and all procedures were approved by the Institutional Review Board at Washington University in St. Louis School of Medicine.

**Table 1:** Patient clinical characteristics.

| Clinical Characteristics | Patient (n=10) |
| --- | --- |
| Severity, n |  |
| Neurapraxia | 6 |
| Axonotmesis | 3 |
| Neurotmesis | 2 |
| # Surgeries (for this injury) | 0.91 ± 0.83 |
| Time since last surgery (mo) | 7.60 ± 12.88 |
| Time since injury (mo) | 9.55 ± 11.94 |
| Affected nerve, n |  |
| Ulnar | 6 |
| Median | 5 |
| Radial | 2 |
| Other | 3 |
| Injury location, n |  |
| Brachial Plexus | 2 |
| Elbow | 2 |
| Forearm | 2 |
| Wrist | 7 |
| Hand | 2 |
| Nerve transfer surgery = Yes | 2 (18%) |
| **Note:** Totals may not add to 10 due to multiple injuries/nerves/locations per patient. |  |

Participants were recruited based on physician referral, medical record search, community recruitment flyers and email lists, and individuals who contacted the laboratory from e.g. clinicaltrials.gov. Inclusion criteria for all participants were: age ≥ 18, English speaking and reading, MRI compatible, and right handed as determined by self report and Edinburgh Handedness Inventory score ≥ +40 (Oldfield, 1971). Additional inclusion criteria for patients were: a chronic (≥ 6 month) peripheral nerve injury (neuropathy with specific localized cause) to the hand, arm, or shoulder; and some impairment to writing, defined as both of: (A) score 2+ (Mild+) on the question “How much difficulty have you had in the last week with writing?” from Disabilities of the Arm, Shoulder, and Hand survey (Beaton et al., 2001); and (B) Box and Blocks motor performance ≥ 1 standard deviation below the mean of age-matched healthy adults (Mathiowetz et al., 1985). For all participants, exclusion criteria were: motor function diagnosis affecting RH in past 2 years (or for patients: diagnoses currently affecting RH unrelated to PNI), motor function diagnosis affecting LH in past 2 years, upper extremity surgery in past 2 months, concussion in past 6 months, amputation (affecting any part of thumb, index, or middle fingers), current chemotherapy, or history of PNI-unrelated conditions that affect brain structure or connectivity (e.g. chronic pain, chronic cocaine use, stroke, traumatic brain injury).

Sample size was based on a power analysis using preliminary data (N = 26) (Kim et al., 2023), which showed potential positive hand_LH-RH_ effects (i.e. LH > RH) in BOLD magnitude in one region of interest (ROI): superior parietal lobule ipsilateral to movement, SPL_hIP3_-ipsi (see *ROI Selection* below) (Mumford, 2012). We conducted a simulation-based power analysis using a linear mixed-effects model with a random intercept for each subject. Fixed effects included behavioral covariates (velocity and direction, modeled separately for each hand or as differences or ratios) and demographic covariates (sex, age, and group). Using the preliminary data to approximate the underlying data-generating mechanism, 500 resampled datasets were generated and the planned analysis was applied to simulated samples of increasing size. Under the effect sizes and variability observed in the pilot data, the results suggested that a sample size of approximately N = 30 would provide 90% power to detect the target effect at a two-sided significance level of α = 0.05. As with all pilot-based power calculations, these estimates are conditional on the observed effect sizes and variance structure and should be interpreted as approximate. To accommodate potential participant exclusion we increased the target sample size to N = 39, which led, after exclusions, to the final sample of N = 33 (23 healthy controls and 10 patients with PNI) as described in the first paragraph of this section.

### 2.2 Behavioral Data

The current study was not designed to identify quantitative effects of performance, so performance during the drawing task was used only for exploratory analyses (see below, 2.7.4. *Magnitude ROI Analysis*).

To assess differences in upper limb function without relying on injury-specific measures that would only be available for patients (e.g. injury severity, time since surgery), all participants self-reported their current upper limb function via the Disabilities of the Arm, Shoulder, and Hand (DASH), a 30-item measure of upper-extremity disability (Beaton et al., 2001; Hudak et al., 1996). The DASH is designed to assess participants’ difficulty performing daily tasks, symptoms such as pain, tingling, weakness, and difficulty sleeping over the past week; and the impact of injury on social activities, work, and self-esteem. For healthy adults, 3/30 DASH items must be omitted (impact of injury on social activities, work, and self-esteem), but nevertheless in the general population the DASH has been established to have high validity and reliability (Hunsaker et al., 2002). Here, to set positive values as reflecting better function, the outcome measure “ability” was calculated as: *ability = 100% – disability%*.

### 2.3 fMRI Acquisitions

Scans were performed on a Siemens PRISMA 3T MRI scanner. fMRI BOLD EPIs were collected using a T2*-weighted gradient echo sequence, a standard 64-channel birdcage radio-frequency coil, and the following parameters: TR = 662 ms, TE = 30 ms, flip angle = 52°, 72 x 72 voxel matrix, FOV = 216 mm, 60 contiguous axial slices acquired in interleaved order, resolution: 3.0 x 3.0 x 3.0 mm, bandwidth = 2670 Hz/pixel, multi-band acceleration = 6x. Siemens auto-align was run at the start of each session. High-resolution T1-weighted structural images were also acquired, using the 3D MP-RAGE pulse sequence: TR = 4500 ms, TE = 3.16 ms, TI = 1000 ms, flip angle = 8.0°, 256 x 256 voxel matrix, FOV = 256 mm, 176 contiguous axial slices, resolution: 1.0 x 1.0 x 1.0 mm. A T2-weighted image was also acquired at: TR = 3000 ms, TE = 409 ms, 256 x 256 voxel matrix, FOV = 256 mm, 176 contiguous axial slices, resolution: 1.0 x 1.0 x 1.0 mm. Spin echo field maps were also collected before the functional runs.

### 2.4 fMRI Task

Each participant used an MRI-compatible tablet (**Figure 1A**) to complete six BOLD functional scans while performing a visually guided precision drawing task based on the STEGA app and previous studies of precision drawing in fMRI (Philip & Frey, 2014; Philip et al., 2023).

**Figure 1:**
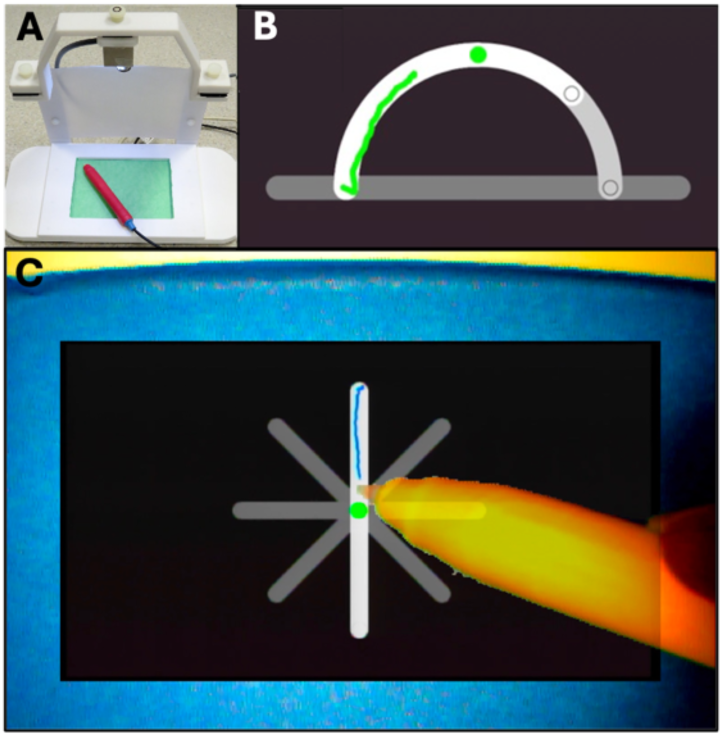
fMRI task. A: MRI-compatible tablet. B: STEGA precision drawing task. C: Participant view of bluescreen integration.

Participants performed the task using one hand per scan, alternating between left and right hand across the six scans (three scans per hand). During the task, participants saw hollow shapes, and were instructed to draw a line within the bounds of the shape (**Figure 1B**), as fast as possible while prioritizing staying in-bounds over speed. To minimize head and shoulder motion, participants were also instructed to draw with only movements of their fingers and hand, while keeping their upper arm and forearm as still as possible. Participants received real-time visual feedback via a transparent overlay of hand/pen over the shape and drawn line (**Figure 1C**), provided via the tablet’s bluescreen technology (Karimpoor et al., 2015; Tam et al., 2011). Pen position data were collected at 60 Hz. The task was presented in a block design with 15.2 seconds (23 images) of drawing, followed by 15.2 seconds (23 images) of rest (fixation cross). A scan comprised ten cycles of draw/rest, with an additional rest block at the start, and 3.3 seconds (5 images) of additional rest after the final rest block, leading to a total duration of 5:23 (488 volumes total: 230 drawing, 258 rest) per scan. Immediately prior to the scan, participants completed 1 scan with each hand in a mock fMRI scanner to familiarize themselves with the task and apparatus, and to practice drawing while keeping their arm still.

### 2.5 fMRI Preprocessing

The data were preprocessed using *fMRIPrep* 24.1.1 (Esteban et al., 2019; RRID:SCR_016216), which is based on *Nipype* 1.8.6 (Gorgolewski et al., 2011; RRID:SCR_002502). Many internal operations of *fMRIPrep* use *Nilearn* 0.10.4 (Abraham et al., 2014; RRID:SCR_001362), mostly within the functional processing workflow. Anatomical preprocessing included intensity-nonuniformity correction, skull stripping, tissue segmentation, FreeSurfer surface reconstruction with T2-weighted refinement of the pial surface, and nonlinear normalization to the symmetric MNI152NLin2009cSym template at 2-mm isotropic resolution. For each of the six BOLD runs, head motion was estimated relative to a reference image, susceptibility distortions were corrected using a B0 field map derived from two EPI reference images, and the BOLD reference was coregistered to the T1-weighted image using six-degree-of-freedom boundary-based registration. Motion-correction, distortion-correction, coregistration, and normalization transforms were concatenated and applied in a single resampling step. fMRIPrep also estimated motion- and tissue-based confounds, including framewise displacement and anatomical and temporal CompCor components. The complete fMRIPrep methods description and software details are provided in **Supplementary Methods S.1.1.1-S.1.1.3**.

### 2.6 fMRI Motion Outlier Identification and Screening

Three methods were used to identify and exclude participants based on motion, task-correlated motion, or poor task compliance. This process evaluated the 37 participants who completed the task, and led to the exclusion of four participants (total) who are therefore not included in the final sample (n = 33).

Throughout this process, motion was defined as framewise displacement (FD), calculated at each volume using FSL_motion_outliers (Jenkinson et al., 2012).

To screen participants with high motion, two screening criteria were employed: 1) Participant-level FD threshold: Participants were excluded from analysis if their participant-mean FD was > 0.5 mm (Power et al., 2012; Power et al., 2014); based on this criterion, 1 participant was excluded. 2) Within-participant motion outliers: a participant would be excluded if >20% of their data in any scan had FD > 75^th^ percentile + (1.5 * inter-quartile range); no participants met that criterion. For additional details see **Supplementary Methods S.1.4.1**.

To screen participants with excess task-correlated motion, the coefficient of determination (Pearson r^2^) was calculated between FD and the task timecourse. Participants were excluded from analysis if their correlation value exceeded the outlier threshold of 75^th^ percentile + (1.5 * inter-quartile range) for either the timecourse or its derivative. Based on this criterion, 2 participants were excluded. For additional details see **Supplementary Methods S.1.4.2**.

To screen participants who failed to comply with the task, experimenters qualitatively evaluated participant compliance during data collection (a yes/no warning flag); 7 participants were flagged, and explored using 3 objective measures of task compliance (tablet misuse, time not drawing, poor performance). Of the 7 flagged participants, 1 was excluded due to tablet misuse, 3 had already been excluded due to previous criteria, and the other 3 were retained. For additional details see **Supplementary Methods S.1.4.3**.

### 2.7 fMRI Analyses

The task fMRI data were analyzed to identify where BOLD magnitude (“magnitude” for brevity) and/or FC were stronger for LH drawing than RH drawing (hand_LH-RH_). In brief, data were reoriented into hemispheres “ipsilateral/contralateral” (to movement), hand_LH-RH_ effects on magnitude were identified via a generalized linear mixed-effects model (GLME) in each region of interest (ROI), whole-brain magnitude was explored via a GLM analysis performed with FSL 6.0.7.12 (FMRIB, Oxford UK) (Smith et al., 2004), and hand_LH-RH_ modulation of FC was identified via generalized psychophysiological interaction (gPPI) (McLaren et al., 2012) performed with CONN 22.v2407 (Nieto-Castanon, 2020).

#### 2.7.1 Data Reorientation

To allow direct comparison of LH vs. RH movement, the data from RH runs were left-right flipped after motion outlier identification and fMRIPrep preprocessing, including registration to the symmetric template MNI152NLin2009cSym. This produced images where the left side of the image consistently represented the hemisphere ipsilateral to movement, and the right side the hemisphere contralateral to movement. Thereafter, all data were analyzed as “ipsilateral/contralateral” to movement.

#### 2.7.2 Magnitude Analysis (GLM)

For BOLD magnitude analysis via GLM, the reoriented data were integrated from fMRIPrep to FSL using established workarounds (Mumford, 2017), followed by spatial smoothing with a 6 mm FWHM SUSAN filter and high pass temporal filtering with 60 sec cutoff; no other preprocessing was applied via FSL. Explanatory variables (EVs) were modeled, along with their temporal derivatives, according to the block design described above (*section 2.4, fMRI Task*), with additional confound EVs based on head motion parameters (3 variables each for translation and rotation) and motion outliers calculated as described above (*section* 2*.C, fMRI Motion Outlier*). The hemodynamic response was accounted for by convolving the model with a double-gamma function. Run-level contrasts of parameter estimates (COPEs) were calculated for Task vs. Rest. The run-level COPEs (Task > Rest) were averaged across runs using a fixed-effects model to produce participant-level results, which were used for the ROI analyses described in *sections 2.7.3-4* and the whole-brain analyses described in *section 2.7.5*.

#### 2.7.3 Region of Interest (ROI) selection

Twelve ROIs were used, 6 areas in each hemisphere (ipsilateral “-ipsi” and contralateral “-contra” to movement). All ROIs were 4-mm-radius spheres around a central seed voxel; thus at the 2-mm isotropic preprocessed resolution, each ROI contained 33 voxels. The 12 ROIs were selected based on previous work implicating 6 areas’ roles in lateralized control of precision hand movement, as shown in **Table 2**.

**Table 2:** Regions of interest (ROIs)

| **Table 2: Regions of interest (ROIs)** |  |  |
| --- | --- | --- |
| **ROI Label** | **Julich-Brain anatomical assignment** | **Coordinates** |
| M1-ipsi / -contra | M1-3b | ±40, -24, 56 |
| IPS-ipsi / -contra | IPS-hIP2 | ±52, -42, 44 |
| SPL<sub>7A</sub>-ipsi / -contra | SPL-7A | ±20, -62, 68 |
| SPL<sub>hIP3</sub>-ipsi / -contra | SPL-hIP3 | ±34, -58, 46 |
| PMd-ipsi / -contra | PMd-6d3 | ±26, -8, 52 |
| SMA-ipsi / -contra | SMA-6mp | ±6, -6, 56 |
| **Note:** Coordinates in MNI152NLin2009cSym template. ROI labels include sub-areas only when needed for disambiguation. |  |  |

The ROIs’ seeds were determined from the following prior studies and updated here with precise sub-areas from the Julich-Brain v3.0.3 parcellation (Amunts et al., 2020; Dickscheid et al., 2026). The M1 ROI seed (in M1_3b_) was based on the normative sensorimotor hand area (Philip & Frey, 2014; Smith & Frey, 2011). The IPS ROI seed (in IPS-hIP2; human intraparietal area 2) was chosen as the voxel with peak activation from left hemisphere areas that previously showed interhemispheric changes related to learning—specifically, where training-related increases in FC with right M1 correlated with NPH skill learning (Philip & Frey, 2016; fig. 4a). The SPL_7A_ ROI seed was chosen from the left hemisphere areas that previously showed interhemispheric changes related to learning and retention—specifically, where training-related changes in FC with right M1 predicted long-term retention of NPH skill (Philip & Frey, 2016; fig. 5a); for this ROI, the seed was selected as the first local maximum with >50% odds of being in cortex (which was also the local maximum with the greatest odds of being inside the brain, 78%) instead of the voxel of peak activation, because the peak was on the cortical surface (−34, -54, 67) and would have produced an ROI substantially outside the brain. The SPL_hIP3_ ROI seed (human intraparietal area 3) was chosen as the voxel with peak activation from right hemisphere areas that previously showed convergent interhemispheric changes that predicted future learning— specifically, where pre-training FC from both the left M1 and left IPS ROIs correlated with subsequent

**Figure 2:**
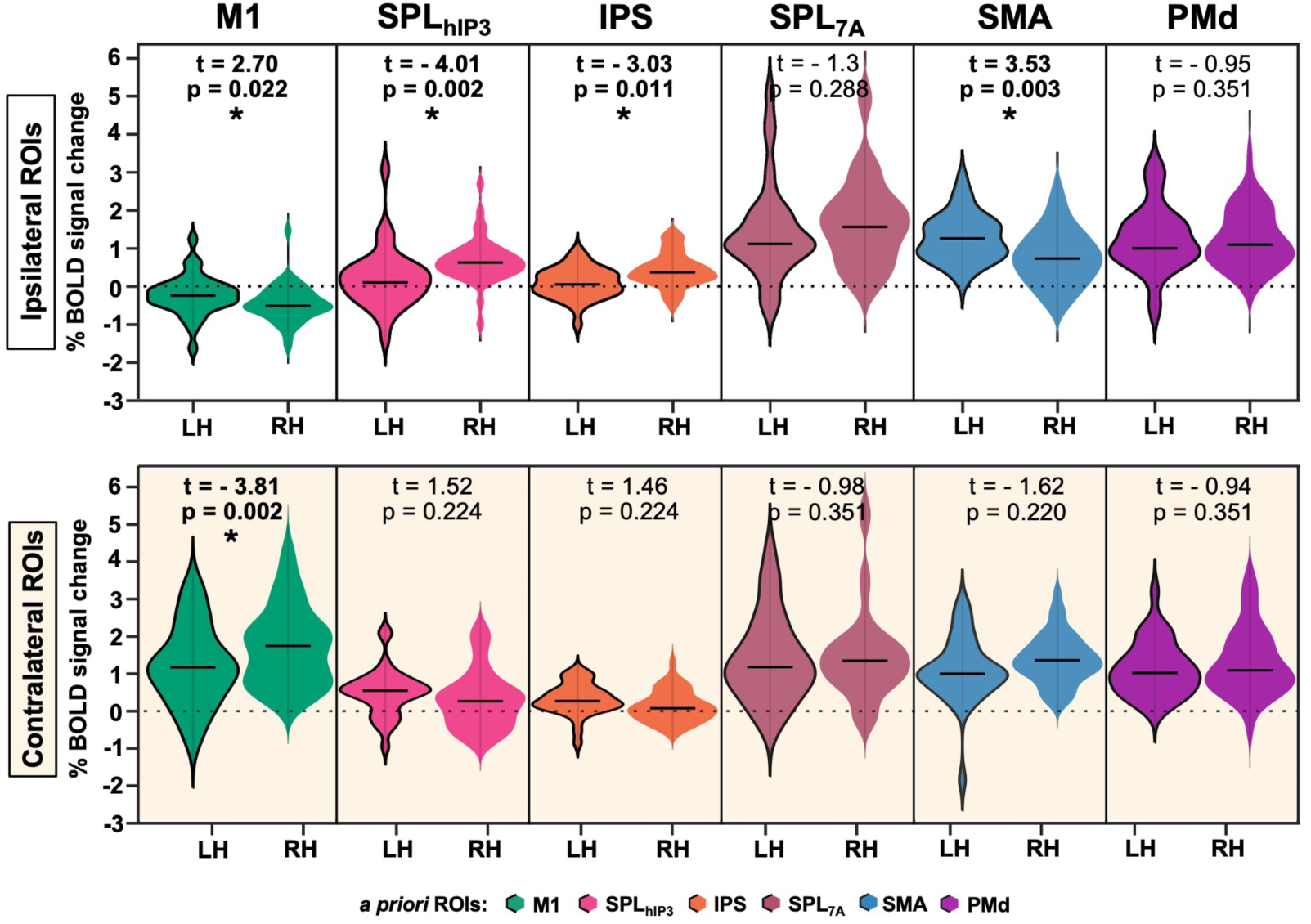
HandLH-RH eBects on BOLD magnitude in a priori ROIs. HandLH-RH eBects were positive in ipsilateral M1 and SMA, and negative in contralateral M1 and ipsilateral SPLhIP3 and IPS. Statistics from GLME.* = p-FDR < 0.05.

**Figure 3.**
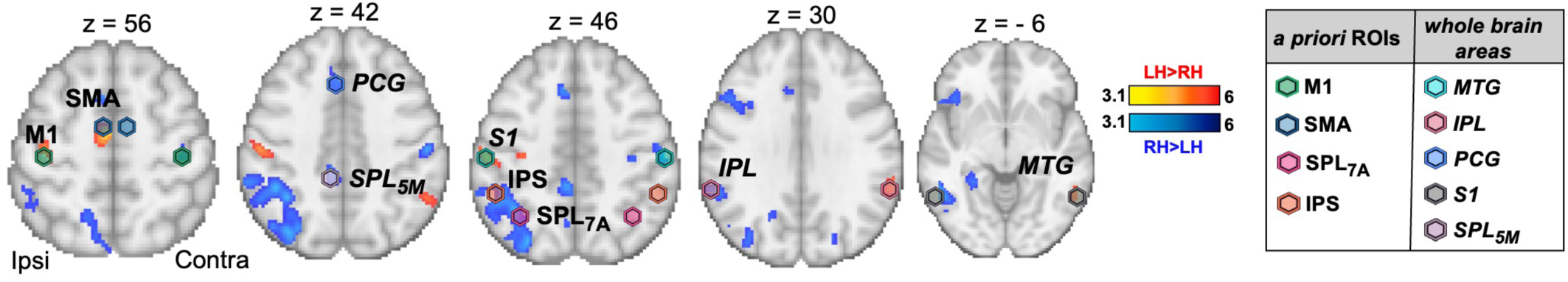
Exploratory whole-brain magnitude eBects, handLH–RH. Results included 7 positive and 12 negative clusters, of which 5 were interpretable (non-overlapping, non-collinear; for selection logic and full anatomical assignments, see Tables 3-4): MTG-contra, IPL-contra, PCG-ipsi, S1-ipsi, and SPL5M-ipsi. A priori ROIs are also shown when they appear on the same slices.

**Figure 4:**
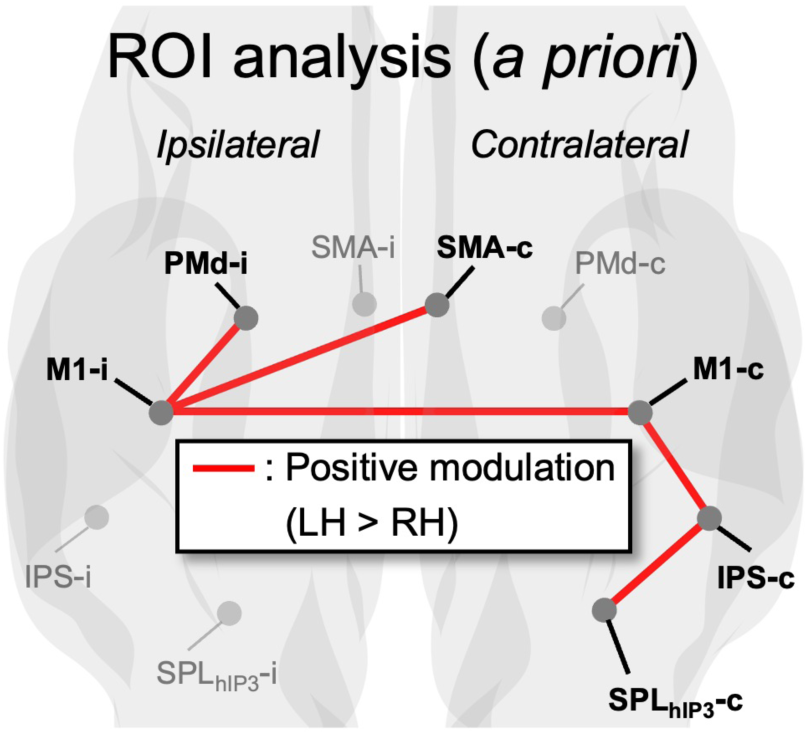
ROI-to-ROI gPPI modulation of functional connectivity (FC), handLH-RH. No connections had negative modulation.

**Figure 5:**
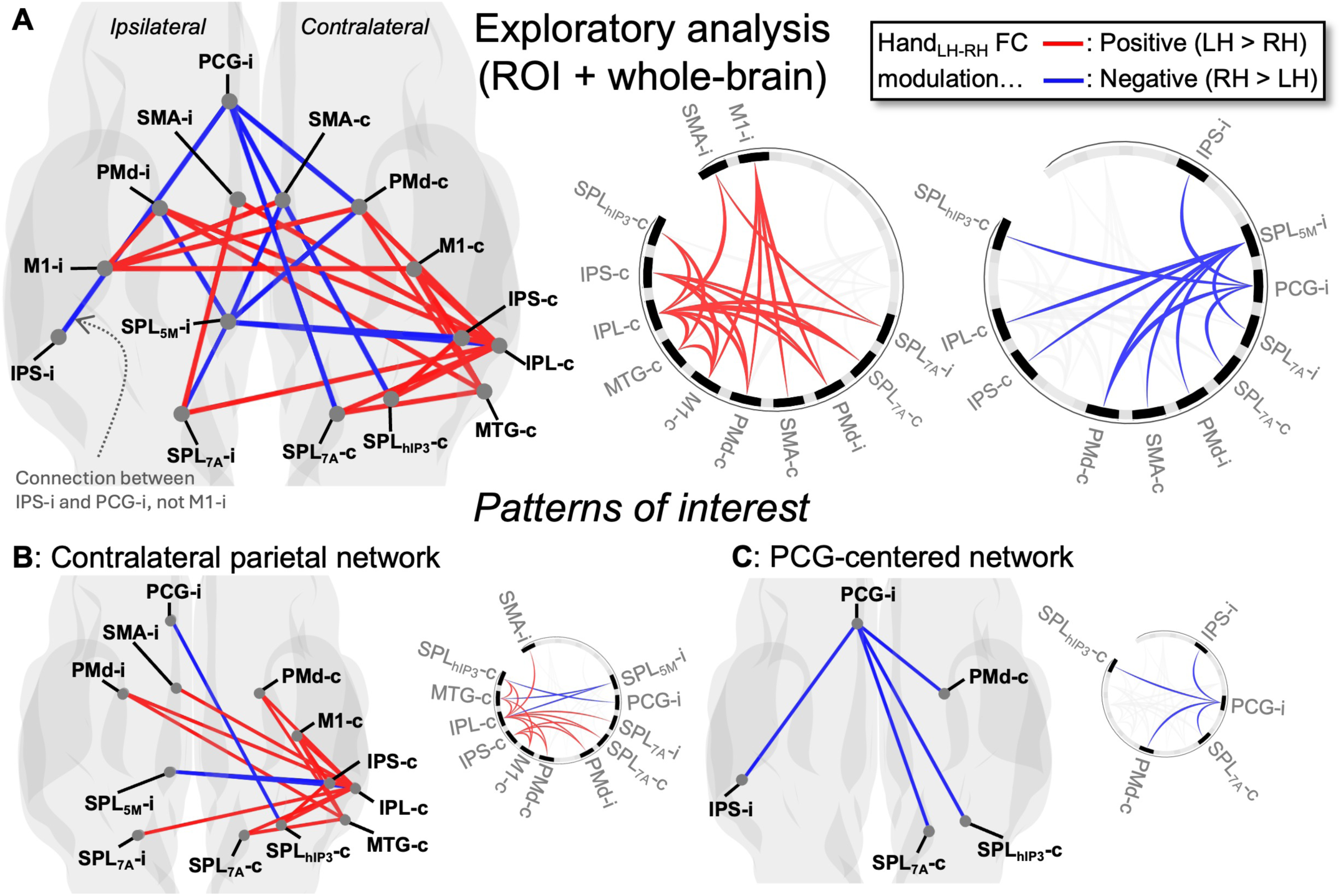
Exploratory analysis of gPPI modulation of functional connectivity (FC), handLH-RH; including 17 areas (12 ROIs + 5 whole-brain areas). Left: glass brain displays. Right: ring plots. A: All connections with significant handLH-RH FC modulation, including two patterns of interest: B: Contralateral parietal network with positive handLH-RH modulation, and C: PCG-centered subnetwork with negative handLH-RH modulation.

NPH learning (Philip et al., 2021; fig. 4). The PMd ROI seed (in PMd-6d3) was chosen as the voxel with peak drawing-specific magnitude in left PMd (Philip et al., 2021; Potgieser et al., 2015). The SMA ROI seed (in SMA-6mp) was chosen as the voxel with peak activation during a hand movement task that previously identified SMA as having a primary role in effective connectivity underpinning lateralized performance (Pool et al., 2014). Spheres were limited to the volume whose chance of being in the appropriate anatomical area according to the Juelich Histological Atlas (Eickhoff et al., 2006; Eickhoff et al., 2007) exceeded a cutoff; the cutoff for parietal and SMA ROIs was ≥ 25%, and PMd was ≥10% because its peak voxel only had a 21% chance of being in Brodmann’s Area 6. However, these anatomical cutoffs did not lead to the removal of any voxels. Seeds from prior studies were transformed to the symmetric template MNI152NLin2009cSym (Fonov et al., 2009) via FLIRT before sphere creation. For brevity, the 12 ROIs are referred to throughout the text and figures using the six labels listed in **Table 2**, which include sub-areas only when needed for disambiguation: M1, IPS, SPL_7A_, SPL_hIP3_, PMd, and SMA.

#### 2.7.4 Magnitude ROI Analysis

To identify whether drawing hand (hand_LH_ vs. hand_RH_) and group (patient, healthy) influenced neural activity in our ROIs, BOLD magnitude in each ROI was modeled using a multivariable generalized linear mixed-effects model (GLME) approach, implemented via the MATLAB function ‘fitglme’ (MATLAB 24.2.0, Mathworks, Natick MA). GLME was chosen because it captures the repeated measures structure of the data (hand_LH_ and hand_RH_ data for each participant) and is suitable for normally distributed data such as our BOLD magnitudes (**Supplementary Figure 1**).

BOLD magnitude from the contrast Task > Rest was converted into “% signal change” using FSL Featquery (Mumford, 2007). Participant-level BOLD magnitudes were calculated as the mean of the 3 runs, separately for each task condition (drawLH or drawRH).

One GLME was created for each of 12 ROIs. In each GLME, participant-level BOLD magnitude was modeled with the repeated-measures factor Participant and 6 factors. The 6 factors were: drawing hand (“hand_LH-RH_,” i.e. LH-drawing > RH-drawing), group (patient, healthy), the interaction of hand_LH-RH_ * group, and 3 participant characteristics (age, sex, and ability).

To assess statistical significance of individual GLME factors, multiple comparisons were controlled via Benjamini–Hochberg false discovery rate (FDR) (Benjamini & Hochberg, 1995) with significance set at FDR-corrected p < 0.05. FDR correction was performed once for the primary hypothesis (hand_LH-RH_; 12 values, one per GLME), secondary hypothesis (group and hand_LH-RH_*group; 24 values, 2 per GLME), and exploratory factors (participant characteristics; 36 values, 3 per GLME). The effect size of Group (patient vs. healthy adult) was calculated for each ROI as Cohen’s *d* via MATLAB function ‘meanEffectSize’.

To assess statistical significance of each GLME model vs. chance, the likelihood ratio test (MATLAB ‘compare’) was used. “Chance” was determined via a bootstrap method. In each draw, factor values were randomly permuted across participants, independently for each factor; and a p-value was calculated using the likelihood ratio test. Reported p-values represent the average across 500 draws.

To explore possible effects of drawing performance on BOLD magnitude, exploratory GLMEs were created that assessed the role of performance confounds and more broadly explored potential relationships between drawing performance and BOLD magnitude. These exploratory GLMEs found no significant effects of any performance variable, and thus are described in **Supplementary Text S.1.7-8**.

#### 2.7.5 Exploratory whole-brain magnitude analysis

Whole-brain analyses were performed to explore the data and identify additional areas with hand_LH-RH_ effects to use as post hoc ROIs. Whole-brain analyses were performed using a mixed-effects model across participants (i.e. at the group level), including participant-specific EVs to capture the repeated-measures structure of LH vs. RH data within each participant, and one EV of interest. For the primary analysis of whole-brain magnitude data, the EV of interest was hand_LH-RH_ (i.e. LH-RH differences within participants), coded as +1 for LH data and –1 for RH data. Secondary analyses were performed with alternate EVs of interest: group and hand_LH-RH_ * group interaction. During all whole-brain analyses, Z-statistic images were created using cluster thresholding at Z > 3.1, p < 0.05.

To describe clusters, coordinates were reported for the center of gravity (CoG) unless the CoG was outside the cluster (e.g. a concave cluster), in which case the coordinates were reported for the peak voxel. To provide anatomical assignments for cluster CoGs, the clusters were warped from symmetric MNI152NLin2009cSym space to FSL asymmetric space with FSL FLIRT, and these asymmetric-space clusters were compared to the Jülich-Brain v3.0.3 parcellation (Amunts et al., 2020; Eickhoff et al., 2007) or MNI152 Cerebellar Atlas; if no assignment with probability ≥10% existed in either atlas, or the Jülich assignment was a GapMap (uncharted cortical area), the Harvard-Oxford Cortical Structural atlas was used. Asymmetric-space clusters were used only for this anatomical assignment process.

To constrain interpretation of exploratory fMRI results, clusters were not retained for further interpretation if they met any of the following criteria: (1) their anatomical assignment was “outside study scope,” i.e. not a parietal, frontal, or temporal cortical area with a potential role in execution of upper limb movement; (2) their volume overlapped with any ROI; or (3) their timecourse was collinear with an *a priori* ROI or with another cluster that met the preceding criteria. Collinearity was defined as Pearson’s |*r*| > 0.7 (Dormann et al., 2012); if two clusters were collinear, the one with higher Z-max was retained for interpretation.

#### 2.7.6 Functional Connectivity (FC) G Generalized Psychophysiological Interaction (gPPI) analysis

To identify hand-dependent (hand_LH-RH_) modulation of FC, a gPPI analysis was performed in the CONN toolbox (22.v2407). First, the reoriented data were smoothed using spatial convolution with a Gaussian kernel of 6 mm FWHM; no other preprocessing was applied via CONN. Denoising was performed using the standard CONN pipeline (Nieto-Castanon, 2020), which included regressing out confounding signals from white matter and CSF BOLD signals (five components each), based on the anatomical component-based noise correction method (aCompCor) (Behzadi et al., 2007) along with six realignment parameters and their first derivatives, outlier scans, condition effects and their first-order derivatives, and linear detrending. High-pass filtering was subsequently applied to retain signal fluctuations above 0.008 Hz.

ROI-to-ROI functional connectivity modulation (hand_LH-RH_) between the twelve ROIs was evaluated via gPPI. For each seed–target ROI pair, the subject-level model included three predictors: (i) HRF-convolved psychological regressors for condition (drawRH, drawLH, and rest); (ii) the physiological regressor (the seed ROI BOLD time series); and (iii) condition-specific interaction terms (PPI) formed by multiplying (i) and (ii), indexing modulation of connectivity between the ROIs. For each connection, the condition-dependent modulation was quantified as the bivariate regression coefficient (β) for the PPI (i.e. interaction) term. At the group level, these PPI β values were analyzed with connection-wise random-effects GLMs across participants to test task effects; the primary contrast was drawLH > drawRH (i.e. hand_LH-RH_). Multiple comparisons were controlled using CONN’s parametric, connection-based framework with Benjamini–Hochberg false discovery rate (FDR) across connections; significance was set at FDR-corrected p < 0.05.

An additional post-hoc exploratory gPPI analysis was performed to explore FC modulation outside our *a priori* ROIs. For each whole-brain cluster retained for interpretation in the exploratory magnitude analysis (*section 2.7.5)*, the cluster’s central voxel was used to create a 4-mm-radius spherical ROI, matching the size of the *a priori* ROIs. To distinguish these new whole-brain-cluster ROIs from *a priori* ROIs, these whole-brain-cluster ROIs are hereafter referred to as “whole-brain areas” instead of “ROIs.” The exploratory gPPI analysis therefore included 17 areas: 12 *a priori* ROIs and 5 whole-brain areas.

Two additional FC analyses were performed. First, to contextualize gPPI’s connectivity-modulation values with a measurement of actual connectivity, standard ROI-to-ROI FC (not gPPI) was estimated separately for LH > rest and RH > rest (e.g. Agren & Hoppe, 2024). Second, the influence of group (patient vs. healthy adult) on gPPI was assessed by examining effects of group and the group*hand_LH-RH_ interaction.

#### 2.7.7 fMRI data quality assessment

Data quality was evaluated using two approaches: BOLD magnitude quality via task-effect signal to noise ratio (SNR), and FC quality via quality control-functional connectivity (QC-FC) analyses. In SNR analysis, each participant’s SNR was calculated as σ (Task>Rest) / σ (model) and averaged across runs (Masharipov et al., 2024), using σ calculated by FSL. The QC-FC analyses involved two processes, both implemented in CONN (Morfini et al., 2023; Nieto-Castanon, 2025). First, the effectiveness of FC denoising was assessed by correlating FC estimates for each connection with participants’ mean FD, and the resulting QC–FC correlation distribution was compared with a permutation-derived null distribution. Second, distance-dependent motion effects were evaluated by regressing QC–FC correlations on the anatomical distance between connected regions and calculating the proportion of variance explained by distance (r²).

## 3. RESULTS

During unimanual execution of a visually guided precision drawing task, M1 contralateral to movement showed a LH-specific combination of lower BOLD magnitude and greater functional connectivity (FC) with two networks: M1/premotor ipsilateral to movement, and posterior parietal areas contralateral to movement. Exploratory analyses identified an additional RH-specific network centered on ipsilateral paracingulate gyrus (PCG).

None of these effects differed significantly between healthy adults and patients with chronic RH peripheral nerve injury (PNI). All results include all 33 participants (23 healthy controls and 10 patients with peripheral nerve injury).

### 3.1 *Drawing with the LH* involves higher BOLD magnitude in ipsilateral frontal motor and contralateral parietal regions of interest

For our primary analysis, we tested whether BOLD magnitude in our 12 *a priori* ROIs (6 areas * 2 hemispheres, ipsilateral and contralateral to movement) varied with drawing hand (i.e. hand_LH-RH_) or other factors (age, sex, DASH upper limb ability, group, and the interaction of group * hand_LH-RH_). For participant-level factor data, see **Supplementary Table 2**.

We found significant effects of hand_LH-RH_ on BOLD magnitude in 5/12 ROIs, but no effects involving other factors. We visualized and quantified the hand_LH-RH_ effects in **Figure 2**; in brief, hand_LH-RH_ was positively associated with BOLD magnitude in M1-ipsi and SMA-ipsi, but hand_LH-RH_ was negatively associated with BOLD magnitude in M1-contra, SPL_hIP3_-ipsi, and IPS-ipsi. No other factors approached significance (p-uncorrected > 0.12; for full model details see **Supplementary Table 3**).

### 3.2 Exploratory whole-brain analysis reveals LH-specific BOLD signal in sensorimotor and parieto-temporal areas

To explore effects of drawing hand on BOLD magnitude across the brain, we performed an exploratory whole-brain group-level analysis of hand_LH-RH_. These whole-brain results (**Figure 3**) included 7 clusters with positive hand_LH-RH_ effects (**Table 3**), divided between sensorimotor regions ipsilateral to movement (M1-6d1, SMA-6mp, S1-BA2, putamen) and parieto-temporal regions contralateral to movement (inferior parietal lobule, IPL-PFm; and middle temporal gyrus, MTG-GapMap, in an “uncharted part of the temporal-to-parietal lobes”) (Amunts et al., 2020). We also found 12 clusters with negative hand_LH-RH_ effects (**Table 4**), including ipsilateral parieto-fronto-temporal regions (IPL-PFm, SPL-7A, SPL-5M, parahippocampal gyrus Ph2, fusiform gyrus FG4, paracingulate gyrus PCG-GapMap (within an “uncharted part of the frontal lobe” on medial surface) (Amunts et al., 2020)), two white matter clusters, contralateral sensory regions (S1-BA2, V3a-hOc4d), and bilateral cerebellar regions (VI ipsilateral, Crus I contralateral).

**Table 3:** Significant clusters in whole-brain BOLD contrast LH>RH (hand_LH-RH_).

| Cluster Index | Voxels | p | Z-MAX | Center of Gravity x, y, z (mm) | Anatomical assignment | Interpreted? (If yes, whole-brain area label) |
| --- | --- | --- | --- | --- | --- | --- |
| 7 | 421 | <0.001 | 4.85 | -28, -24, 68 * | M1-6d1 (ipsi) | No, ROI overlap |
| 6 | 204 | 0.001 | 5.36 | -6.9, -9.6, 59 * | SMA-6mp (ipsi) | No, ROI overlap |
| 5 | 136 | 0.013 | 5.46 | 61.6, -43.3, 30.5 | IPL-PFm (contra) | Yes (IPL-contra)<sup>1</sup> |
| 4 | 125 | 0.019 | 4.76 | -54.0, -21.1, 44.4 | S1-BA2 (ipsi) | Yes (S1-ipsi) |
| 3 | 121 | 0.021 | 4.21 | -26.7, -2.1, 5.28 | Putamen (ipsi) | No, scope |
| 2 | 99 | 0.047 | 4.35 | 55.7, -54.2, -7.0 | MTG-GapMap (contra) | Yes (MTG-contra) |
| 1 | 98 | 0.049 | 3.8 | 53.7, -54.1, 39.1 | IPL-PGa (contra) | No, collinear<sup>1</sup> |

**Notes:** Significance determined at cluster-corrected $p < 0.05$ . Coordinates in MNI152Nlin2009cSym.\* = Concave cluster, coordinates represent peak instead of CoG. “Scope” = not a cortical area involved in upper limb movement execution.<sup>1</sup> = Clusters #1 and #5 were collinear ( $r = 0.812$ ), so cluster #1 was not interpreted.

**Table 4:** Significant clusters in whole-brain BOLD contrast RH>LH (negative hand_LH-RH_)

|  |  |  |  |  |  |  |
| --- | --- | --- | --- | --- | --- | --- |
| Cluster Index | Voxels | p | Z-MAX | Center of Gravity x, y, z (mm) | Anatomical assignment | Interpreted? (if yes, whole-brain area label) |
| 12 | 1865 | 6x10<sup>-17</sup> | 5.68 | -66, -38, 18 * | IPL-PFm (ipsi) | No, collinear<sup>1</sup> |
| 11 | 1157 | 3x10<sup>-12</sup> | 4.92 | -44, 38, 8 * | WM (ipsi) | No, scope |
| 10 | 506 | 7x10<sup>-7</sup> | 5.64 | -17.2, -64.2, -19.7 | Cblm VI (ipsi) | No, scope |
| 9 | 367 | 2x10<sup>-5</sup> | 5.21 | -47.1, -59.7, -8.8 | FG4 (ipsi) | No, scope |
| 8 | 329 | 5x10<sup>-5</sup> | 5.94 | -19.3, -56.2, 17.2 | WM callosal body (ipsi) | No, scope |
| 7 | 218 | 0.001 | 5.00 | 56, -20, 44 * | S1-BA2 (contra) | No, collinear<sup>2</sup> |
| 6 | 188 | 0.002 | 4.82 | -6.7, 19.8, 43.2 | PCG-GapMap (ipsi) | Yes (PCG-ipsi) |
| 5 | 182 | 0.003 | 4.33 | -29.8, -40.9, -11.5 | Ph2 (ipsi) | No, scope |
| 4 | 175 | 0.003 | 4.35 | -10.5, -63.2, 54.8 | SPL-7A (ipsi) | No, ROI overlap |
| 3 | 139 | 0.012 | 4.68 | -6.5, -38.7, 43.2 | SPL-5M (ipsi) | Yes (SPL<sub>5M</sub>-ipsi) |
| 2 | 138 | 0.012 | 5.22 | 32.2, -64.7, -27.7 | Cblm Crus I (contra) | No, scope |
| 1 | 101 | 0.044 | 5.12 | 26.9, -86.7, 24.4 | V3a-hOc4d (contra) | No, scope |

**Notes:** Significance determined at cluster-corrected p < 0.05. Coordinates in MNI152NLin2009cSym. Cblm = cerebellum, WM = white matter.\* and “scope” defined in Table 3.<sup>1</sup> = Collinear with ROIs in SPL<sub>hip3</sub>-ipsi (r = 0.829) and IPS-ipsi (r = 0.775).<sup>2</sup> = Collinear with ROI in M1-contra (r = 0.829).

Before interpreting whole-brain results, we constrained our interpretation of exploratory results in fMRI data by excluding clusters that were assigned to anatomical regions outside the study scope (upper limb execution), overlapped with ROIs, or were collinear with other clusters (**Supplementary Table 4**). This led to 5 interpretable new clusters from the whole-brain analysis: positive hand_LH-RH_ effects in IPL-PFm, S1-BA2, and MTG-GapMap (**Table 3**); and negative hand_LH-RH_ effects in PCG-GapMap and SPL-5M (**Table 4**).

Hereafter, these five clusters and the corresponding cluster-derived spheres used in subsequent analyses are referred to as the whole-brain areas IPL-contra (from IPL-PFm), S1-ipsi (from S1-BA2), MTG-contra (from MTG-GapMap), PCG-ipsi (from PCG-GapMap), and SPL_5M_-ipsi (from SPL-5M).

### 3.3 Functional connectivity: LH drawing involves increased connectivity between contralateral motor cortex and bilateral parieto-motor networks

We evaluated hand-specific modulation of FC between the 12 ROIs via a generalized psychophysiological interaction (gPPI) analysis, which revealed significant hand_LH-RH_ modulation in 5 connections between 8/12 ROIs, as listed in **Table 5** and shown in **Figure 4**.

**Table 5:** Connections with significant handLH-RH modulation.

| ROI 1 | ROI 2 | T-score | p-uncorrected | p-FDR |
| --- | --- | --- | --- | --- |
| PMd-ipsi | M1-ipsi | 4.37 | 1.21x10<sup>-4</sup> | 0.0160 |
| M1-contralateral | M1-ipsi | 4.00 | 3.48x10<sup>-4</sup> | 0.0230 |
| IPS-contralateral | SPL<sub>hip3</sub>-contralateral | 3.80 | 6.19x10<sup>-4</sup> | 0.0272 |
| M1-contralateral | IPS-contralateral | 3.50 | 0.0014 | 0.0453 |
| M1-ipsi | SMA-contralateral | 3.41 | 0.0018 | 0.0470 |

To contextualize this hand-dependent modulation, we examined traditional FC (not gPPI modulation) between ROIs, separately for each drawing condition (LH, RH). All connections identified by gPPI had FC > 0 in both conditions, with numerically stronger FC during LH (t-score range 6.31– 12.64, FDR-corrected p < .001) than RH (t-score range 2.63– 7.47, FDR-corrected p ≤ .015); for details see **Supplementary Table 5**. Notably, the strongest connection during LH drawing was between PMd-ipsi and PMd-contra; this connection was also second-strongest during RH ROIs + the 5 interpretable whole-brain areas identified above drawing. Overall, these results indicate that the positive gPPI modulation for hand_LH-RH_ reflects a strengthening of pre-existing positive connectivity. On a post-hoc basis, we expanded the gPPI analysis to 17 areas: the 12 *a priori* (*section 3.2*). This revealed significant hand_LH-RH_ modulation in 44 connections (including the original 5) between 15/17 areas, as shown in **Figure 5A** and detailed in **Supplementary Table 6**. New hand_LH-RH_ patterns included increased FC in contralateral parietal network (**Figure 5B**), and decreased FC between ipsilateral paracingulate gyrus (PCG) and an asymmetric parietal-premotor network (**Figure 5C**).

### 3.4 Magnitude-connectivity integration reveals motor and parietal asymmetries during LH drawing

To integrate our fMRI analyses, we visualized our BOLD magnitude and FC modulation effects together in **Figure 6**. Combining the two fMRI modalities in our *a priori* ROIs (**Figure 6A**) revealed two hand_LH-RH_ effects. First, the corticospinal output area M1-contra had reduced magnitude but increased FC. M1-contra’s increased FC included stronger interhemispheric FC with M1-ipsi, which had higher magnitude and stronger FC with PMd-ipsi and SMA-contra; and stronger intrahemispheric FC with contralateral parietal regions IPS and SPL_hIP3_, which had unchanged magnitude. Second, parietal cortex showed an asymmetric pattern: parietal areas SPL_hIP3_ and IPS showed greater FC in the contralateral hemisphere, but decreased magnitude in the ipsilateral hemisphere.

**Figure 6:**
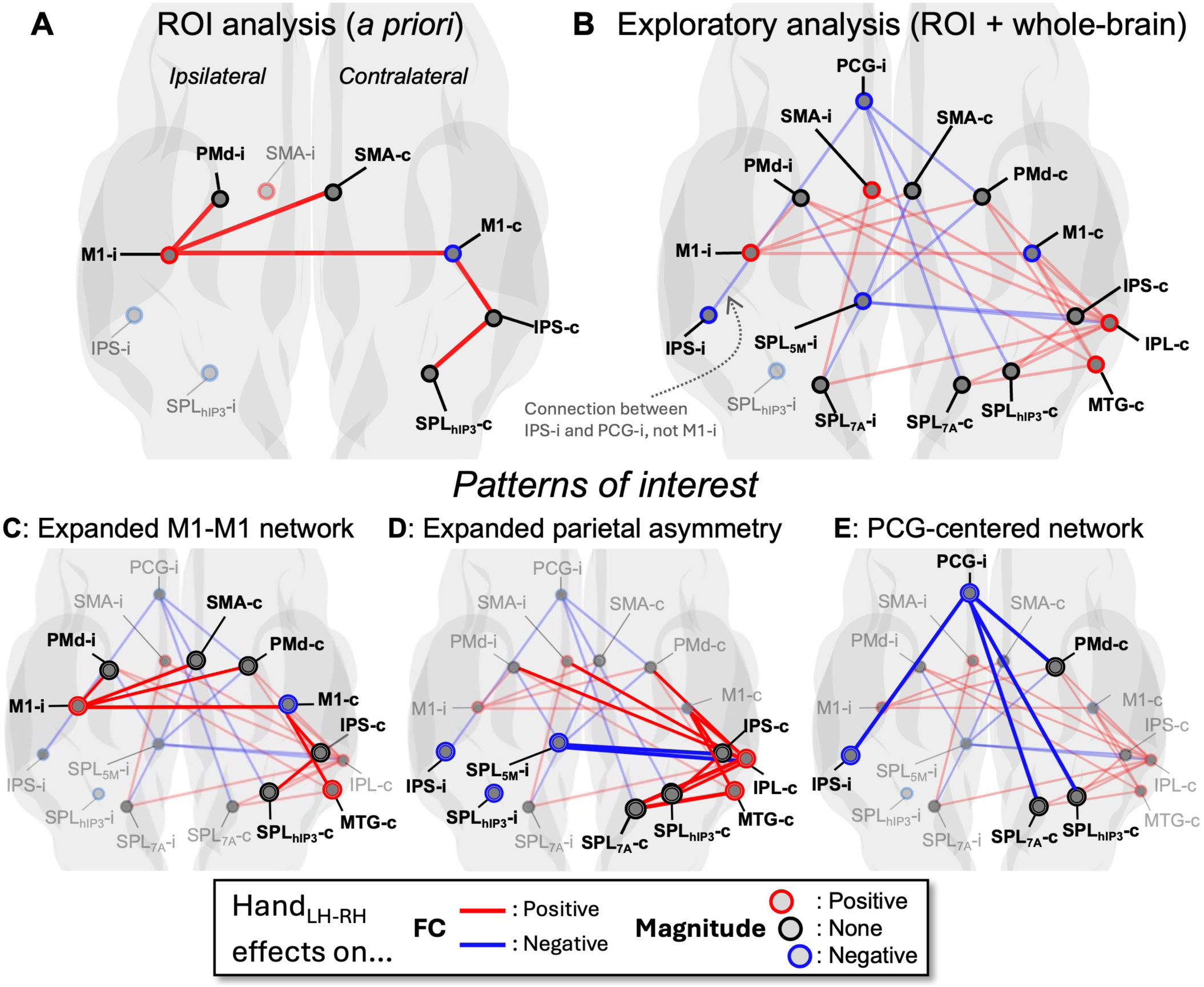
Integrated visualization of BOLD magnitude and gPPI modulation of functional connectivity (FC), handLH-RH. Areas are shown if they had any eBects in magnitude (circles) or FC (lines). A: A priori ROIs. B-E: Exploratory analysis, including a priori ROIs and areas identified from whole-brain exploration. B: All areas, including 3 patterns of interest (C-E): C: Expanded M1–M1 network, D: Expanded parietal asymmetry, and E: Exploratory PCG-centered network.

We found additional hand_LH-RH_ effects in our exploratory analysis (combining *a priori* ROIs and whole-brain areas), as shown in **Figure 6B**. The M1 pattern described in the previous paragraph remained largely unchanged, with two additional connections (**Figure 6C**): M1-contra’s intrahemispheric FC additionally included MTG-contra, which had higher magnitude; and M1-ipsi increased FC with PMd-contra in addition to the previously observed PMd-ipsi. The involvement of both PMd regions is consistent with high FC between PMd-ipsi and PMd-contra (*section 3.3*).

The exploratory analysis also extended the parietal asymmetry (**Figure 6D**) beyond the network observed in our *a priori* ROIs. In the ipsilateral hemisphere, the pattern of reduced magnitude expanded to include SPL_5M_-ipsi. In the contralateral hemisphere, the pattern of increased FC expanded beyond IPS and SPL_hIP3_ to include IPL-contra and temporal area MTG-contra, both of which also showed increased magnitude during hand_LH-RH_.

Finally, magnitude-connectivity integration clarified the nature of the exploratory PCG-centered network (**Figure 6E**): during hand_LH-RH_, PCG-ipsi had lower magnitude and lower connectivity with a bilateral fronto-parietal network including 3 contralateral areas with hand-nonspecific magnitude (SPL_hIP3_, SPL_7A_, PMd) and one ipsilateral area with reduced magnitude in hand_LH-RH_ (IPS).

#### 3.4.1 Anatomical interpretation of hand_LH-RH_ effects reveals a left motor-premotor network and right parietal asymmetry

All results above were analyzed and reported as “ipsilateral” and “contralateral” to the moving hand (*section 2.7.1*), to allow direct comparison between LH and RH drawing. Thus, the same ipsilateral or contralateral label corresponds to different anatomical hemispheres depending on the drawing hand: for example, the ipsilateral hemisphere is the left hemisphere during LH drawing, but the right hemisphere during RH drawing. Therefore, converting from ipsilateral/contralateral to left/right can clarify the hemispheric organization of the effects described above.

The hand_LH-RH_ effects in M1-contra (**Figure 6A, C**) represent M1-right having lower magnitude and higher parietal/interhemispheric FC, whereas the M1-ipsi effects represent M1-left having higher magnitude and higher FC with premotor regions. The parietal asymmetry (**Figure 6A, D**) included two hand_LH-RH_ effects: first, increased FC in contralateral parietal regions, which represents greater FC within a right parietal network (and between this network and contralateral right M1); second, negative hand_LH-RH_ magnitude effects primarily in ipsilateral parietal regions, which are equivalent to positive RH effects (i.e. hand_RH-LH_) in ipsilateral parietal regions, and therefore represent greater BOLD magnitude in right parietal regions during RH drawing. Thus, the right-hemisphere parietal network showed increased FC during LH drawing (with increased magnitude in exploratory only, IPL/MTG), and increased magnitude (including 2/4 a priori areas) during RH drawing. Finally, the exploratory PCG network (**Figure 6B, E**) showed negative hand_LH-RH_ effects, which are equivalent to positive RH effects centered on right PCG.

Overall, combining BOLD magnitude and FC modulation identified two hand_LH-RH_ mechanisms during precision drawing. First, right M1 showed lower BOLD magnitude but greater connectivity with a left M1-PMd network and right parieto-temporal areas (IPS, SPL_hIP3_, IPL, and MTG). Second, the right parietal cortex exhibited an asymmetric pattern, showing greater FC during LH drawing but greater magnitude primarily during RH drawing. In addition, exploratory analyses identified a right PCG-centered network with greater magnitude and connectivity during RH drawing.

### 3.5 Peripheral nerve injury did not interact with hand-specific effects

To address our hypothesis that patients with chronic PNI would use the same neural mechanisms as healthy adults for LH drawing, we first evaluated whether Group (patient vs. healthy) or a Group * Hand_LH-RH_ interaction contributed to the GLME of BOLD magnitude in any ROI (or any whole-brain area). We found no significant effects of Group, nor of Group * Hand_LH-RH_ interactions (p-FDR > 0.76), with one non-significant trend toward a Group * Hand_LH-RH_ interaction in whole-brain area MTG-contra (p-uncorrected = 0.0238; all other p-uncorrected > 0.12), as detailed in **Supplementary Table 7**. To confirm this negative result, we evaluated the univariate effect size of Group in each ROI and whole-brain area, and found that Group’s 95% confidence interval always included zero, whether combining data from LH and RH drawing (p > 0.07, **Figure 7**) or separately analyzing LH and RH (p > 0.22). In addition, gPPI analysis revealed no connections that had significant gPPI modulation (positive or negative) with Group or Group*Hand_LH-RH_. Therefore, we found no evidence that any of our hand_LH-RH_ effects differed between patients and healthy adults.

**Figure 7:**
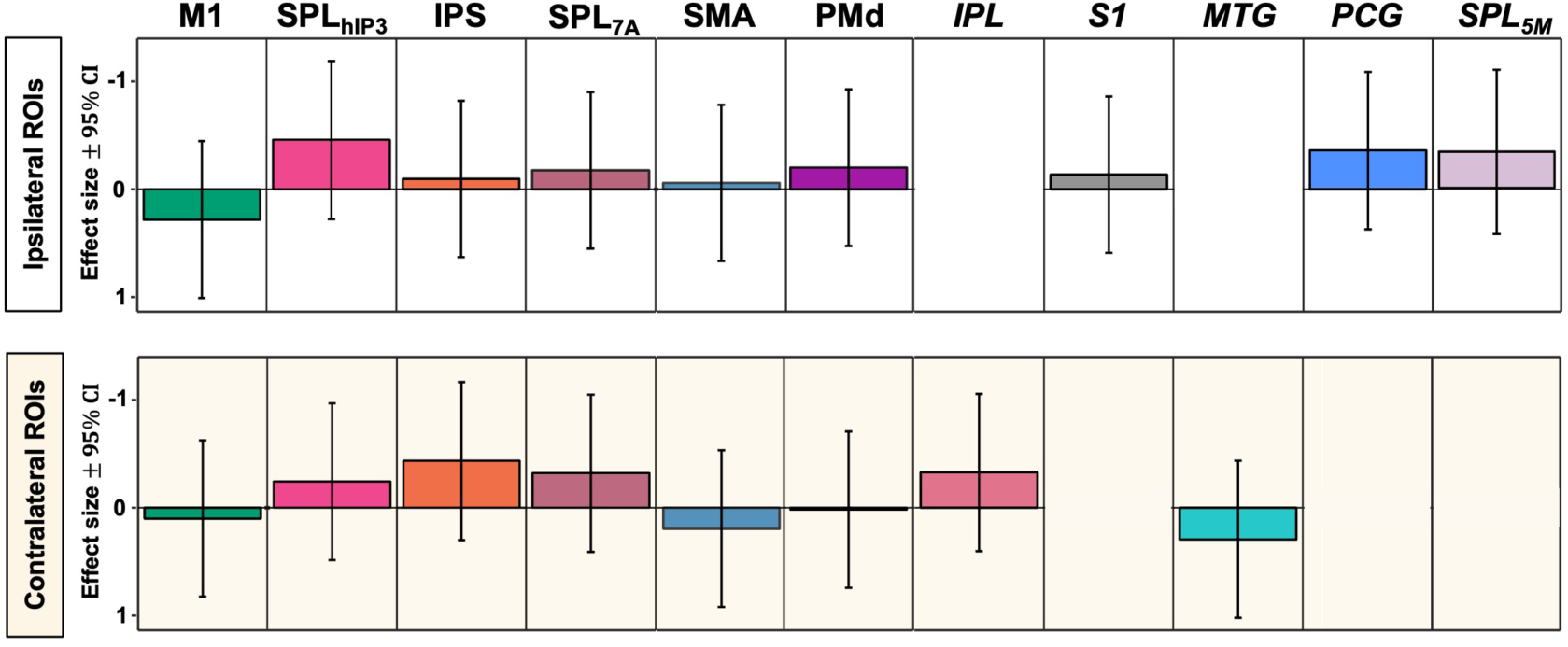
No eBect of group (PNI patients, n =10, vs. healthy controls, n=23) on BOLD magnitude. In every ROI and whole-brain cluster (italics), 95% confidence intervals of eBect size (Cohen’s d) include zero

### 3.6 fMRI data quality assessment

To quantify BOLD magnitude data quality, the SNR of the task effect (Draw > Rest) was 0.487 ± 0.08. To quantify FC data quality after denoising, the mean QC–FC was 0.01 (SD = 0.18), and the QC-FC distribution overlapped 97.4% with the permutation-derived null distribution (compared with 95.9% before denoising), consistent with minimal residual motion-related associations. Connection distance explained less than 0.1% of the variance in QC–FC correlations (R² < .001), indicating negligible linear distance dependence.

## 4. DISCUSSION

Our study used a visually guided precision drawing task to characterize neural activity and functional connectivity associated with left hand (LH) drawing in right-handed individuals, with and without unilateral peripheral nerve injury (PNI) to the right hand (RH). We found three hand-dependent mechanisms, two *a priori* and one exploratory (**Figure 8**): 1) a left hemisphere motor-premotor network that supported bilateral precision control through hand-dependent patterns of intra- and inter-hemispheric interactions; 2) a right hemisphere parietal network associated with drawing with either hand, but differently for each hand; and 3) an exploratory paracingulate gyrus (PCG)-centered network of bilateral fronto-parietal areas, associated with RH drawing. These neural mechanisms did not differ between healthy adults and patients with chronic RH PNI.

**Figure 8:**
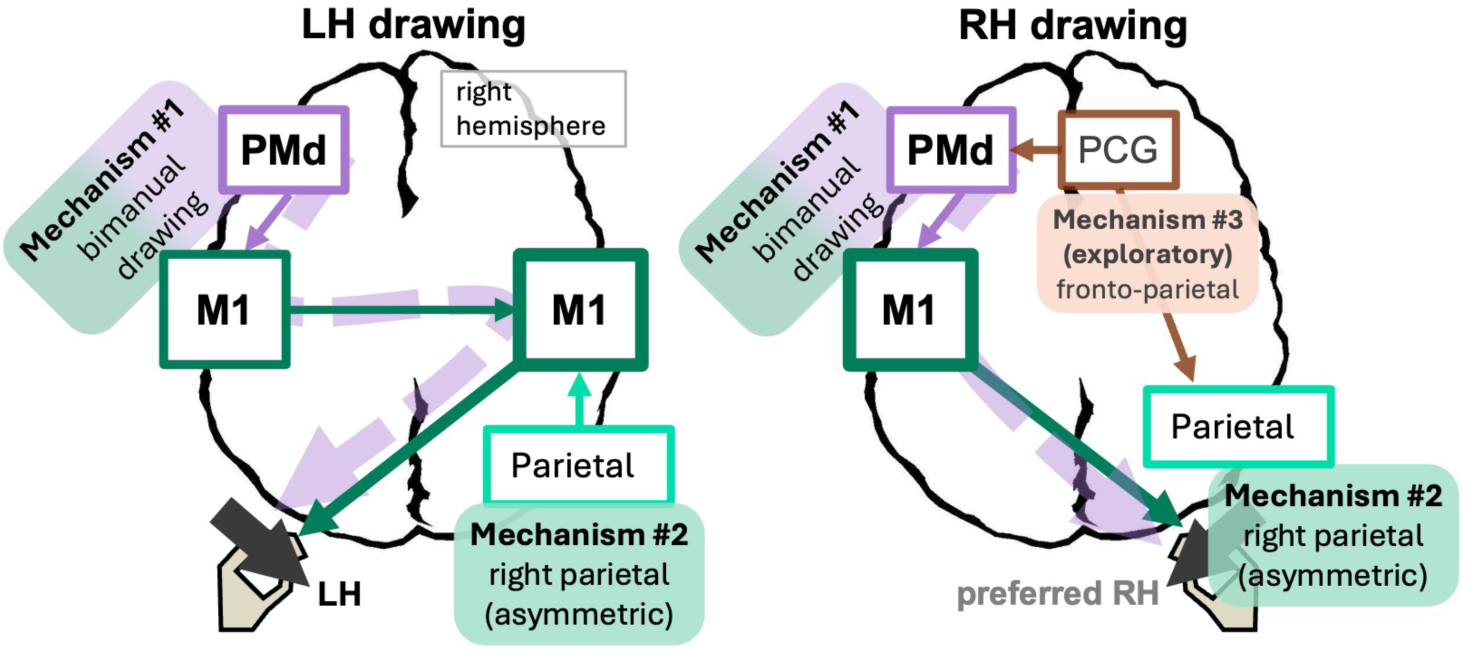
Model of neural mechanisms with digerential involvement in left hand (LH) vs. right hand (RH) precision drawing, in right-handed adults. Mechanism #1 is a left M1-PMd network for bimanual drawing, with access to contralateral M1 via hand-specific paths (dashed lines): an indirect interhemispheric path during LH drawing, and a direct intrahemispheric path during RH drawing. Mechanism #2 is a righthemisphere parietal network involved digerently in LH vs. RH drawing, with greater connectivity (internal and parietal-motor) during LH drawing. Mechanism #3 (exploratory) is a fronto-parietal network centered on right paracingulate gyrus (PCG), involved preferentially in RH drawing.

### 4.1 Left-hemisphere motor-premotor network supports bimanual precision drawing

We found a left-lateralized primary motor-dorsal premotor (M1-PMd) network that functions as a shared neural mechanism for bimanual precision drawing in right-handed adults. This network may involve M1’s role in ipsilateral motor control under complex or high-demand conditions (Barany et al., 2020; Buetefisch et al., 2014; Kim et al., 1993; Verstynen et al., 2005) and left PMd’s role in the temporal aspects of bilateral movement and their associated task-specific demands (Kornysheva & Diedrichsen, 2014; Schluter et al., 1998; Yang et al., 2024). During RH drawing, this network includes contralateral left M1 as the source of descending corticospinal output. During LH drawing, the same network remains engaged, but control of the acting hand is mediated through contralateral right M1 via interhemispheric M1-M1 connectivity. Thus, the left-lateralized M1-PMd network retains its role across hands by adaptively recruiting contralateral motor regions through interhemispheric M1-M1 connections (**Figure 8**, Mechanism #1).

This left-lateralized M1-PMd network provides anatomical specificity to existing models of hand dominance, which propose that RH advantages in right-handed individuals arise from a left hemisphere specialization for predictive dynamics that enable precise movement (complementary dominance), and LH disadvantages for precise movement arise from interhemispheric delays (Kitchen et al., 2025; Liepmann, 1905; Shabbott,B.A., & Sainburg, R.L., 2008). It is also consistent with previous findings that FC in a M1-PMd-somatosensory network supports precision manual tasks with the RH (Yang et al., 2024). Here we extend this work by identifying a specific left-lateralized mechanism and demonstrating that its access to corticospinal motor output area differs across hands: direct intrahemispheric during RH drawing, and indirect interhemispheric during LH drawing.

Our effector-independent role for left PMd diverges from the findings of Potgieser et al. (2015), who reported bilateral PMd activation during a visually guided figure-copying task. This difference may depend on task characteristics: unlike Potgieser et al. (2015), our task imposed constraints on spatial precision, which affects lateralization across parietal and premotor cortices (Haaland et al., 2004).

### 4.2 Right parietal network is asymmetrically engaged during LH and RH drawing

We identified a right-lateralized parietal network with an asymmetric hand-dependent role: greater FC within the network (and with contralateral right M1) during LH drawing, but greater magnitude during RH drawing. This asymmetry was observed in our *a priori* ROIs (intraparietal sulcus human intraparietal area 2, IPS; and superior parietal lobule human intraparietal area 3, SPL_hIP3_), and extended in our exploratory analysis to a parieto-temporal network (superior parietal lobule area 5 medial, SPL_5M_-ipsi; inferior parietal lobule IPL-contra; and middle temporal gyrus MTG-contra). This asymmetric pattern suggests that the right parietal network contributed to both LH and RH drawing through distinct neural mechanisms: stronger distributed parieto-motor integration during LH drawing, and stronger regional recruitment during RH drawing (**Figure 8**, Mechanism #2).

This right parietal network is unlikely to represent the Complementary Dominance Hypothesis’ right hemisphere specialization for impedance-related aspects of movement (Kitchen et al., 2025), because such a specialization should show increased interhemispheric connectivity during RH drawing (when a right hemisphere mechanism supports left M1), whereas we found increased parietal-motor FC during LH drawing only. The specific role of this right parietal network in LH drawing cannot yet be specified because right parietal cortex is involved in numerous visuospatial, sensorimotor, and goal-related processes during precision hand control (Cavina-Pratesi et al., 2018; Tunik et al., 2008). These potential roles include motor skill learning (Della-Maggiore et al., 2004), action planning with the contralateral hand (Valyear & Frey, 2015), and aspects of visuomotor control that depend on higher-order goals, such as goal-directed visuospatial attention (Corbetta et al., 2000; Wang et al., 2016), goal-movement integration (Averbeck et al., 2009; Wolpert et al., 1998), intended sensory outcomes (Krasovsky et al., 2014), and evaluation of action consequences (Hamilton & Grafton, 2006, 2008). Our right parietal network also extended into MTG (though we retained the label “parietal” since MTG was an exploratory result), which has a similar range of involvement in goal-directed movement: e.g. monitoring intersensory conflict (van Kemenade et al., 2019) and visual evaluation of action outcomes (Jastorff et al., 2010).

To specify the role of this right parietal network in LH control, future studies with a larger sample size are required to link the network’s activity/connectivity to specific behavioral performance characteristics.

Nevertheless, our findings demonstrate that the network is recruited differently during RH and LH drawing. The greater magnitudes observed during RH drawing in right IPS, IPL and (exploratory) SPL_5M_ may reflect stronger localized recruitment of sensorimotor integration systems during RH precision control. In contrast, LH drawing did not affect magnitude in our *a priori* ROIs (increased magnitude in exploratory right MTG and IPL, which might reflect increased action-monitoring requirements); instead, LH drawing involved increased FC throughout the right parietal network and with output area M1-contra, suggesting greater reliance on the parietal network’s distributed visuomotor coordination during LH control.

### 4.3 Exploratory analyses identified an RH-specific right paracingulate network

Our exploratory analyses identified a bilateral parieto-frontal network associated with RH precision drawing, centered on ipsilateral right paracingulate gyrus (PCG). During RH drawing, right PCG showed both greater BOLD magnitude and stronger FC with left IPS as well as right PMd, SPL_hIP3_, and SPL_7A_. Unlike the right parietal findings above, none of this network’s areas or connections showed LH-specific increases in FC or magnitude, nor did we identify any effects in contralateral left PCG, suggesting that this PCG-centered parieto-frontal network was specific to RH drawing (**Figure 8**, Mechanism #3).

The functional significance of this PCG finding remains uncertain. Our PCG locus is in a medial frontal “GapMap” region within the Julich-Brain atlas, which indicates a portion of cortex that remains incompletely defined cytoarchitectonically and functionally (Amunts et al., 2020). Nevertheless, the observed connectivity pattern suggests potential involvement in higher-order coordination between frontal motor and parietal visuomotor systems during RH drawing. However, this PCG-centered finding was exploratory and requires replication in larger studies.

### 4.4 Proposed hand-dependent mechanisms may remain accessible after chronic peripheral nerve injury

We found no evidence of differences between patients with chronic RH PNI and healthy adult controls, in the three hand-specific mechanisms listed above (*sections 4.1-4.3*). Although the modest patient sample size (23 healthy adults vs. 10 patients) limits definitive conclusions regarding group equivalence, these findings suggest that chronic unilateral PNI may not impede use of the dominant-hemisphere motor-premotor network, along with its associated interhemispheric and parieto-motor coordination networks. The potential preservation of these networks after PNI may carry important clinical implications, because these networks could represent potential targets for neuromodulatory or behavioral interventions aimed at improving function of the non-preferred hand (NPH). Multiple rehabilitation approaches have demonstrated efficacy in enhancing NPH performance, including bimanual training (Stinear & Byblow, 2004), mirror therapy (Dohle et al., 2009; Nojima et al., 2012), action observation (Ertelt et al., 2007), motor imagery (Sharma et al., 2006), and bihemispheric stimulation strategies such as dual tDCS or rTMS (Lindenberg et al., 2010; Waters-Metenier et al., 2014). These approaches converge on the principle that bilateral cortical recruitment, particularly via the dominant hemisphere, can facilitate motor recovery or compensation after unilateral impairment.

In addition, this work helped establish a preliminary normative template of bilateral cortical control for precision movement in the healthy brain. Such a template may ultimately prove useful for understanding and improving rehabilitation of asymmetric neurological conditions such as stroke, wherein some patients must rely on their NPH due to chronic PH impairment. For example, future studies could determine whether patients recruit typical cortical control strategies (e.g., the left M1-PMd network identified here) or rely on alternative compensatory mechanisms, and whether altered interhemispheric connectivity contributes to variability in NPH performance. Such approaches may help characterize atypical neural mechanisms associated with unilateral motor impairment and inform development of individualized, mechanism-driven rehabilitation strategies.

### 4.5 Ipsilateral superior parietal lobule may play a learning-dependent role

Our results did not support our initial hypothesis that our locus in ipsilateral SPL human intraparietal area 3 (SPL_hIP3_) would play a greater role during LH than RH drawing. SPL_hIP3_ did show recruitment for LH drawing (increased FC, unchanged magnitude), but in contralateral rather than ipsilateral SPL_hIP3_; in other words, right SPL_hIP3_ is part of our right parietal network (*section 4.2*) that is more connected to M1 motor output during LH drawing, compared to left SPL_hIP3_’s connection during RH drawing. We based our initial hypothesis on previous findings that interhemispheric FC involving ipsilateral left SPL_hIP3_ could predict future LH skill learning (Philip et al., 2021). However, the current study did not involve learning: participants completed only 5.5 minutes of familiarization with each hand, followed by 16.5 minutes with each hand during fMRI, far below the amount of time required for right-handed adults to learn a LH precision drawing task (4.6 ± 2.3 sessions of 20 minutes each; Philip & Frey, 2016). Therefore, our SPL_hIP3_ locus may have captured an ipsilateral mechanism specific to LH skill learning.

## 5. Limitations

This study had a modest sample size (n = 33). However, this sample size was appropriate for detecting our primary task-related activation effects (i.e., hand_LH-RH_ contrasts), as determined by our *a priori* power analysis. Further supporting our modest sample size, our drawing movement task produced larger neural effects than the designs used to create most sample size benchmarks, as demonstrated by our SNR of 0.49, approaching the 0.5 practical upper limit for task fMRI (Masharipov et al., 2024). Nevertheless, we adapted our analysis to our modest sample size by declining to evaluate the quantitative brain-behavior relationships that are most likely to produce unstable results at small sample sizes (Grady et al., 2020), and treating gPPI as a secondary analysis because simulation studies suggest that gPPI effects in a block design study of SNR ∼0.5 require n ≥ 50 (Masharipov et al., 2024). For quantitative analyses (e.g. performance) and robust gPPI results, further studies with larger sample sizes are necessary [link to pre-registration will be added after peer review]. Our findings involving whole-brain areas, including IPL, MTG, PCG, and SPL_5M_, were also exploratory and identified after the primary ROI analyses. Mechanism #3 relies entirely on these exploratory results, and thus should be considered preliminary and hypothesis-generating until replicated in larger, independently powered datasets. In addition, the patient subgroup was small, limiting power to detect subtle group differences or injury-specific adaptations. Larger studies will be needed for robust brain-behavior analyses, stronger gPPI inference, and more definitive patient-control comparisons.

Our results describe neural mechanisms specific to the control of precision drawing (a PH-specialized task) in right-handed adults. Therefore, our findings provide no information about neural mechanisms of NPH specializations, nor about left-handed individuals. Moreover, functional brain lateralization can vary among people with the same handedness (Kroliczak et al., 2021). Rehabilitation strategies should therefore consider individual functional organization rather than handedness alone.

## 6. Conclusion

This study identified two primary neural mechanisms that distinguish LH and RH precision drawing in right-handed adults. First, a left-lateralized M1-PMd network that supported bilateral drawing; this network engaged interhemispherically during LH drawing and intrahemispherically during RH drawing. Second, a right parietal network which was engaged during bilateral drawing, but differently for each hand, with increased functional connectivity (within-network and parietal-motor) during LH drawing and greater regional recruitment during RH drawing. In addition, exploratory analyses suggested that RH drawing may engage a parieto-premotor network centered on right paracingulate gyrus. Together, these results suggest that RH and LH control may rely on different modes of visuomotor network engagement, including differential engagement of effector-independent systems in left premotor and right parietal cortices.

Participants with chronic RH PNI did not show detectable differences from healthy adults in these proposed mechanisms, although the modest patient sample limits conclusions about group equivalence. Overall, these findings suggest that precision hand control depends on flexible interactions among dominant-hemisphere motor-premotor systems, interhemispheric motor pathways, and distributed visuomotor coordination networks. The left M1-PMd network, in particular, may represent a candidate target for future neuromodulatory or behavioral interventions aimed at improving non-preferred hand function after unilateral injury.

## Data and Code Availability

The data that supports the findings of this study are openly available at OpenNeuro, https://doi.org/10.18112/openneuro.ds008162.v1.0.3 (Kapil et al., 2026). The codes that support the findings of this study will be made openly available upon completion of clinical trial NCT05207878.

## Author Contributions

NK: software, formal analysis, investigation, data curation, writing-original draft, writing-review & editing, visualization. TK: methodology, software, investigation, data curation, writing-original draft, writing- review C editing. SG: investigation, writing-review & editing. RZ: methodology, formal analysis, writing- review C editing. ARC: conceptualization, writing-review & editing, funding acquisition. IGD: conceptualization, writing-review & editing, funding acquisition. LL: methodology, writing-review & editing, funding acquisition. MPM: software, methodology, writing-review & editing, funding acquisition. MDW: formal analysis, writing-review & editing. YW: conceptualization, writing-review & editing, funding acquisition. DMB: resources, writing-review & editing. CJD: resources, writing-review & editing. SEM: resources, writing-review & editing. BAP: conceptualization, methodology, investigation, writing-original draft, writing-review & editing, visualization, resources, supervision, and funding acquisition.

## Declaration of Competing Interest

Benjamin A. Philip reports financial support was provided by National Institute of Neurological Disorders and Stroke. Washington University School of Medicine and author Benjamin A. Philip have a licensing agreement with PlatformSTL to commercialize a version of the drawing task used in this work. This agreement has not been executed, and no financial or non-financial support has occurred. The other authors declare that they have no known competing financial interests or personal relationships that could have appeared to influence the work reported in this paper.

## Funding

This work was supported by NINDS R01 NS114046 to Benjamin A. Philip. Data storage was funded by NCI grant P30 CA091842 and NCATS grant UL1 TR002345.

## Acknowledgements

The authors would like to thank Summer Fletcher for their help with data collection and logistics, and Joshua Shimony for his help with patient safety. This study was funded by NIH/NINDS grant R01NS114046 to BAP. Data storage was funded by NCI grant P30 CA091842 and NCATS grant UL1 TR002345 to Washington University.

## Notes

https://doi.org/10.18112/openneuro.ds008162.v1.0.3

### Competing Interest Statement

Author BAP has a licensing agreement with PlatformSTL to commercialize a version of the drawing task used in this work.

### Summary of Updates

Increased specificity and clarity in anatomical/ROI labels throughout the manuscript. Added MRI data quality validation. Clarified various methodological details, and moved some to Supplementary.

## References

Abraham, A., Pedregosa, F., Eickenberg, M., Gervais, P., Mueller, A., Kossaifi, J., Gramfort, A., Thirion, B., & Varoquaux, G. (2014). Machine learning for neuroimaging with scikit-learn. Frontiers in neuroinformatics, 8, 14.

Agren, T., & Hoppe, J. M. (2024). Extensive functional connectivity between brain areas implicated in mental imagery production and phobic fear during both emotional and neutral mental imagery. Behav Brain Res, 462, 114893. 10.1016/j.bbr.2024.114893

Amunts, K., Mohlberg, H., Bludau, S., & Zilles, K. (2020). Julich-Brain: A 3D probabilistic atlas of the human brain’s cytoarchitecture. Science, 369(6506), 988–992.

Averbeck, B. B., Crowe, D. A., Chafee, M. V., & Georgopoulos, A. P. (2009). Differential contribution of superior parietal and dorsal-lateral prefrontal cortices in copying. Cortex, 45(3), 432–441. 10.1016/j.cortex.2008.02.007

Barany, D. A., Revill, K. P., Caliban, A., Vernon, I., Shukla, A., Sathian, K., & Buetefisch, C. M. (2020). Primary motor cortical activity during unimanual movements with increasing demand on precision. Journal of Neurophysiology, 124(3), 728–739.

Beaton, D. E., Davis, A. M., Hudak, P., & McConnell, S. (2001). The DASH (Disabilities of the Arm, Shoulder and Hand) outcome measure: what do we know about it now? The British Journal of Hand Therapy, 6(4), 109–118.

Behzadi, Y., Restom, K., Liau, J., & Liu, T. T. (2007). A component based noise correction method (CompCor) for BOLD and perfusion based fMRI. NeuroImage, 37(1), 90–101. 10.1016/j.neuroimage.2007.04.042

Benjamini, Y., & Hochberg, Y. (1995). Controlling the false discovery rate: a practical and powerful approach to multiple testing. Journal of the Royal statistical society: series B (Methodological), 57(1), 289–300.

Blank, R., Miller, V., & Von Voss, H. (2000). Human motor development and hand laterality: a kinematic analysis of drawing movements. Neuroscience Letters, 295(3), 89–92.

Bottenhorn, K. L., Flannery, J. S., Boeving, E. R., Riedel, M. C., Eickhoff, S. B., Sutherland, M. T., & Laird, A. R. (2018). Cooperating yet distinct brain networks engaged during naturalistic paradigms: A meta-analysis of functional MRI results. Network Neuroscience, 3(1), 27–48. 10.1162/netn_a_00050

Buetefisch, C. M., Revill, K. P., Shuster, L., Hines, B., & Parsons, M. (2014). Motor demand-dependent activation of ipsilateral motor cortex. Journal of Neurophysiology, 112(4), 999–1009. 10.1152/jn.00110.2014

Cavina-Pratesi, C., Connolly, J. D., Monaco, S., Figley, T. D., Milner, A. D., Schenk, T., & Culham, J. C. (2018). Human neuroimaging reveals the subcomponents of grasping, reaching and pointing actions. Cortex, 98, 128–148. 10.1016/j.cortex.2017.05.018

Coito, A., Michel, C. M., Vulliemoz, S., & Plomp, G. (2019). Directed functional connections underlying spontaneous brain activity. Hum Brain Mapp, 40(3), 879–888. 10.1002/hbm.24418

Corbetta, M., Kincade, J. M., Ollinger, J. M., McAvoy, M. P., & Shulman, G. L. (2000). Voluntary orienting is dissociated from target detection in human posterior parietal cortex. Nature neuroscience, 3(3), 292–297.

Della-Maggiore, V., Malfait, N., Ostry, D. J., & Paus, T. (2004). Stimulation of the posterior parietal cortex interferes with arm trajectory adjustments during the learning of new dynamics. The Journal of Neuroscience: the Official Journal of the Society for Neuroscience, 24(44), 9971–9976. 10.1523/JNEUROSCI.2833-04.2004

Dexheimer, B., Sainburg, R. L., Sharp, S., & Philip, B. A. (2024). Roles of Handedness and Hemispheric Lateralization: Implications for Rehabilitation of the Central and Peripheral Nervous Systems: A Rapid Review. American Journal of Occupational Therapy, 78(2). 10.5014/ajot.2024.050398

Dickscheid, T., Gui, X., Simsek, A. N., Schiffer, C., Mangin, J.-F., Leprince, Y., Jirsa, V., Bjaalie, J. G., Leergaard, T. B., Bludau, S., & Amunts, K. (2026). Siibra: a software tool suite for realizing a Multilevel Human Brain Atlas from complex data resources. Nature methods, 23(8), 1647–1658. 10.1038/s41592-026-03159-x

Dohle, C., Püllen, J., Nakaten, A., Küst, J., Rietz, C., & Karbe, H. (2009). Mirror Therapy Promotes Recovery From Severe Hemiparesis: A Randomized Controlled Trial. Neurorehabilitation and Neural Repair, 23(3), 209–217. 10.1177/1545968308324786

Dormann, C. F., Elith, J., Bacher, S., Buchmann, C., Carl, G., Carré, G., Marquéz, J. R. G., Gruber, B., Lafourcade, B., Leitão, P. J., Münkemüller, T., McClean, C., Osborne, P. E., Reineking, B., Schröder, B., Skidmore, A. K., Zurell, D., & Lautenbach, S. (2012). Collinearity: a review of methods to deal with it and a simulation study evaluating their performance. Ecography, 36(1), 27–46. 10.1111/j.1600-0587.2012.07348.x

Dyck, P. J., Boes, C. J., Mulder, D., Millikan, C., Windebank, A. J., Dyck, P. J. B., & Espinosa, R. (2005). History of standard scoring, notation, and summation of neuromuscular signs. A current survey and recommendation. Journal of the Peripheral Nervous System, 10(2), 158–173.

Eickhoff, S., Heim, S., Zilles, K., & Amunts, K. (2006). Testing anatomically specified hypotheses in functional imaging using cytoarchitectonic maps. NeuroImage, 32(2), 570–582. 10.1016/j.neuroimage.2006.04.204

Eickhoff, S. B., Paus, T., Caspers, S., Grosbras, M.-H., Evans, A. C., Zilles, K., & Amunts, K. (2007). Assignment of functional activations to probabilistic cytoarchitectonic areas revisited. [Review]. NeuroImage, 36(3), 511–521. 10.1016/j.neuroimage.2007.03.060

Ertelt, D., Small, S., Solodkin, A., Dettmers, C., McNamara, A., Binkofski, F., & Buccino, G. (2007). Action observation has a positive impact on rehabilitation of motor deficits after stroke. NeuroImage, 36, T164–T173. 10.1016/j.neuroimage.2007.03.043

Esteban, O., Markiewicz, C. J., Blair, R. W., Moodie, C. A., Isik, A. I., Erramuzpe, A., Kent, J. D., Goncalves, M., DuPre, E., & Snyder, M. (2019). fMRIPrep: a robust preprocessing pipeline for functional MRI. Nature methods, 16(1), 111–116.

Finn, E. S., Glerean, E., Hasson, U., & Vanderwal, T. (2022). Naturalistic imaging: The use of ecologically valid conditions to study brain function. NeuroImage, 247, 118776. 10.1016/j.neuroimage.2021.118776

Finn, E. S., Glerean, E., Khojandi, A. Y., Nielson, D., Molfese, P. J., Handwerker, D. A., & Bandettini, P. A. (2020). Idiosynchrony: From shared responses to individual differences during naturalistic neuroimaging. NeuroImage, 215, 116828. 10.1016/j.neuroimage.2020.116828

Fonov, V. S., Evans, A. C., McKinstry, R. C., Almli, C. R., & Collins, D. (2009). Unbiased nonlinear average age-appropriate brain templates from birth to adulthood. NeuroImage(47), S102.

Freud, E., Macdonald, S. N., Chen, J., Quinlan, D. J., Goodale, M. A., & Culham, J. C. (2018). Getting a grip on reality: Grasping movements directed to real objects and images rely on dissociable neural representations. Cortex, 98, 34–48. 10.1016/j.cortex.2017.02.020

Fu, Z., Tu, Y., Di, X., Biswal, B. B., Calhoun, V. D., & Zhang, Z. (2017). Associations between Functional Connectivity Dynamics and BOLD Dynamics Are Heterogeneous Across Brain Networks. Front Hum Neurosci, 11, 593. 10.3389/fnhum.2017.00593

Gorgolewski, K., Burns, C. D., Madison, C., Clark, D., Halchenko, Y. O., Waskom, M. L., & Ghosh, S. S. (2011). Nipype: a flexible, lightweight and extensible neuroimaging data processing framework in python. Frontiers in neuroinformatics, 5, 13. 10.3389/fninf.2011.00013

Grady, C. L., Rieck, J. R., Nichol, D., Rodrigue, K. M., & Kennedy, K. M. (2020). Influence of sample size and analytic approach on stability and interpretation of brain-behavior correlations in task-related fMRI data. Human Brain Mapping, 42(1), 204–219. 10.1002/hbm.25217

Gutierrez-Barragan, D., Basson, M. A., Panzeri, S., & Gozzi, A. (2019). Infraslow state fluctuations govern spontaneous fMRI network dynamics. Current Biology, 29(14), 2295–2306. e2295.

Haaland, K. Y., Elsinger, C. L., Mayer, A. R., Durgerian, S., & Rao, S. M. (2004). Motor sequence complexity and performing hand produce differential patterns of hemispheric lateralization. *Cognitive Neuroscience*, Journal of, 16(4), 621–636. http://ieeexplore.ieee.org/xpls/abs_all.jsp?arnumber=6789895

Hamilton, A. F., & Grafton, S. T. (2006). Goal Representation in Human Anterior Intraparietal Sulcus. Journal of Neuroscience, 26(4), 1133–1137. 10.1523/JNEUROSCI.4551-05.2006

Hamilton, A. F., & Grafton, S. T. (2008). Action outcomes are represented in human inferior frontoparietal cortex. Cereb Cortex, 18(5), 1160–1168. 10.1093/cercor/bhm150

Harris, P. A., Taylor, R., Thielke, R., Payne, J., Gonzalez, N., & Conde, J. G. (2009). Research electronic data capture (REDCap)—a metadata-driven methodology and workflow process for providing translational research informatics support. Journal of biomedical informatics, 42(2), 377–381. 10.1016/j.jbi.2008.08.010

Hasson, U., Malach, R., & Heeger, D. J. (2010). Reliability of cortical activity during natural stimulation. Trends in Cognitive Sciences, 14(1), 40–48. 10.1016/j.tics.2009.10.011

Hayashi, M. J., Saito, D. N., Aramaki, Y., Asai, T., Fujibayashi, Y., & Sadato, N. (2008). Hemispheric asymmetry of frequency-dependent suppression in the ipsilateral primary motor cortex during finger movement: a functional magnetic resonance imaging study. Cereb Cortex, 18(12), 2932–2940. 10.1093/cercor/bhn053

He, B., Zhu, Z., Zhu, Q., Zhou, X., Zheng, C., Li, P., Zhu, S., Liu, X., & Zhu, J. (2014). Factors predicting sensory and motor recovery after the repair of upper limb peripheral nerve injuries [Research and Report]. Neural Regeneration Research, 9(6), 661–672. 10.4103/1673-5374.130094

Hudak, P. L., Amadio, P. C., Bombardier, C., Beaton, D., Cole, D., Davis, A., Hawker, G., Katz, J. N., Makela, M., & Marx, R. G. (1996). Development of an upper extremity outcome measure: the DASH (disabilities of the arm, shoulder, and head). American journal of industrial medicine, 29(6), 602–608.

Hunsaker, F. G., Cioffi, D. A., Amadio, P. C., Wright, J. G., & Caughlin, B. (2002). The American academy of orthopaedic surgeons outcomes instruments. J Bone Joint Surg Am, 84(2), 208–215.

Jastorff, J., Clavagnier, S., Gergely, G., & Orban, G. A., (2010). Neural Mechanisms of Understanding Rational Actions: Middle Temporal Gyrus Activation by Contextual Violation. Cerebral Cortex, 21(2), 318–329. 10.1093/cercor/bhq098

Jenkinson, M., Beckmann, C. F., Behrens, T. E. J., Woolrich, M. W., & Smith, S. M. (2012). FSL. *NeuroImage*, C2(2), 782–790. 10.1016/j.neuroimage.2011.09.015

Kapil, N., Kim, T., Gassass, S., Fletcher, S., McAvoy, M. P., & Philip, B. A. (2026). Precision drawing during task fMRI in healthy adults and individuals with peripheral nerve injury. (ds008162; Version 1.0.3) [Dataset]. OpenNeuro. 10.18112/openneuro.ds008162.v1.0.3

Karimpoor, M., Churchill, N. W., Tam, F., Fischer, C. E., Schweizer, T. A., & Graham, S. J. (2018). Functional MRI of Handwriting Tasks: A Study of Healthy Young Adults Interacting with a Novel Touch-Sensitive Tablet. Front Hum Neurosci, 12, 30. 10.3389/fnhum.2018.00030

Karimpoor, M., Tam, F., Strother, S. C., Fischer, C. E., Schweizer, T. A., & Graham, S. J. (2015). A computerized tablet with visual feedback of hand position for functional magnetic resonance imaging. Frontiers in Human Neuroscience, 9, 150.

Kim, S. G., Ashe, J., Hendrich, K., Ellermann, J. M., Merkle, H., Uğurbil, K., & Georgopoulos, A. P. (1993). Functional magnetic resonance imaging of motor cortex: hemispheric asymmetry and handedness. Science, 261(5121), 615–617. 10.1126/science.8342027

Kim, T., Fletcher, S., Gonzalez, C. L. R., & Philip, B. A. (2025). Block Building Task Identifies Distinct Groups of Left/Right-hand Choice Patterns After Unilateral Peripheral Nerve Injury. J Vis Exp(217). 10.3791/66919

Kim, T., Gassass, S., Carter, A. R., Dobbins, I. G., Liu, L., McAvoy, M., Sun, Z., Wang, Y., & Philip, B. A. (2023). Ipsilateral parietal cortex supports precision movement with the non-dominant hand. Progress in Clinical Motor Control II, Chicago, IL.

Kim, T., Lohse, K. R., Mackinnon, S. E., & Philip, B. A. (2024). Patient Outcomes After Peripheral Nerve Injury Depend on Bimanual Dexterity and Preserved Use of the Affected Hand. Neurorehabilitation and Neural Repair, 38(2), 134–147. 10.1177/15459683241227222

Kitchen, N. M., Dexheimer, B., Yuk, J., Maenza, C., Ruelos, P. R., Kim, T., & Sainburg, R. L. (2025). The complementary dominance hypothesis: a model for remediating the ’good’ hand in stroke survivors. Journal of Physiology, 603(3), 663–683. 10.1113/JP285561

Kornysheva, K., & Diedrichsen, J. (2014). Human premotor areas parse sequences into their spatial and temporal features. Elife, 3, e03043. 10.7554/eLife.03043

Krasovsky, A., Gilron, R., Yeshurun, Y., & Mukamel, R. (2014). Differentiating intended sensory outcome from underlying motor actions in the human brain. The Journal of Neuroscience: the Official Journal of the Society for Neuroscience, 34(46), 15446–15454. 10.1523/JNEUROSCI.5435-13.2014

Kringelbach, M. L., Perl, Y. S., Tagliazucchi, E., & Deco, G. (2023). Toward naturalistic neuroscience: Mechanisms underlying the flattening of brain hierarchy in movie-watching compared to rest and task. Science Advances, 9(2), eade6049. 10.1126/sciadv.ade6049

Kroliczak, G., Buchwald, M., Kleka, P., Klichowski, M., Potok, W., Nowik, A. M., Randerath, J., & Piper, B. J. (2021). Manual praxis and language-production networks, and their links to handedness. Cortex, 140, 110–127. 10.1016/j.cortex.2021.03.022

Liepmann, H. M. O. (1905). Die linke Hemisphäre und das Handeln. Münchener Medizinische Wochenschrift. (49), 2322–2326, 2375–2378.

Lindenberg, R., Renga, V., Zhu, L. L., Nair, D., & Schlaug, G. (2010). Bihemispheric brain stimulation facilitates motor recovery in chronic stroke patients. Neurology, 75(24), 2176–2184. 10.1212/wnl.0b013e318202013a

Mani, S., Mutha, P. K., Przybyla, A., Haaland, K. Y., Good, D. C., & Sainburg, R. L. (2013). Contralesional motor deficits after unilateral stroke reflect hemisphere-specific control mechanisms. Brain, 136(Pt 4), 1288–1303. 10.1093/brain/aws283

Marcori, A. J., Monteiro, P. H. M., & Okazaki, V. H. A. (2019). Changing handedness: What can we learn from preference shift studies? Neurosci Biobehav Rev, 107, 313–319. 10.1016/j.neubiorev.2019.09.019

Masharipov, R., Knyazeva, I., Korotkov, A., Cherednichenko, D., & Kireev, M. (2024). Comparison of whole-brain task-modulated functional connectivity methods for fMRI task connectomics. Commun Biol, 7. 10.1101/2024.01.22.576622

Mathiowetz, V., Volland, G., Kashman, N., & Weber, K. (1985). Adult norms for the Box and Block Test of manual dexterity. American Journal of Occupational Therapy, 36(6), 386–391.

McLaren, D. G., Ries, M. L., Xu, G., & Johnson, S. C. (2012). A generalized form of context-dependent psychophysiological interactions (gPPI): a comparison to standard approaches. NeuroImage, 61(4), 1277–1286. 10.1016/j.neuroimage.2012.03.068

Morfini, F., Whitfield-Gabrieli, S., & Nieto-Castanon, A. (2023). Functional connectivity MRI quality control procedures in CONN. Front Neurosci, 17, 1092125. 10.3389/fnins.2023.1092125

Mumford, J. (2007). A guide to calculating percent change with featquery. *Unpublished Tech Report In*: http://mumford.bol.ucla.edu/perchange_guide.pdf, 177.

Mumford, J. A. (2012). A power calculation guide for fMRI studies. Soc Cogn Affect Neurosci, 7(6), 738–742. 10.1093/scan/nss059

Mumford, J. A. (2017). FEAT registration workaround. Retrieved June 1 from https://mumfordbrainstats.tumblr.com/post/166054797696/feat-registration-workaround

Nieto-Castanon, A. (2020). Handbook of functional connectivity Magnetic Resonance Imaging methods in CONN. 10.56441/hilbertpress.2207.6598

Nieto-Castanon, A. (2025). Preparing fMRI data for statistical analysis. In fMRI techniques and protocols (pp. 163–191). Springer.

Nojima, I., Mima, T., Koganemaru, S., Thabit, M. N., Fukuyama, H., & Kawamata, T. (2012). Human Motor Plasticity Induced by Mirror Visual Feedback. The Journal of Neuroscience, 32(4), 1293–1300. 10.1523/jneurosci.5364-11.2012

Oldfield, R. C. (1971). The assessment and analysis of handedness: the Edinburgh inventory. Neuropsychologia, 9(1), 97–113. 10.1016/0028-3932(71)90067-4

Peterson, S. M., Singh, S. H., Wang, N. X. R., Rao, R. P. N., & Brunton, B. W. (2021). Behavioral and Neural Variability of Naturalistic Arm Movements. eneuro, 8(3). 10.1523/ENEURO.0007-21.2021

Philip, B. A., & Frey, S. H. (2014). Compensatory Changes Accompanying Chronic Forced Use of the Nondominant Hand by Unilateral Amputees. The Journal of Neuroscience: the Official Journal of the Society for Neuroscience, 34(10), 3622–3631. 10.1523/JNEUROSCI.3770-13.2014

Philip, B. A., & Frey, S. H. (2016). Increased functional connectivity between cortical hand areas and praxis network associated with training-related improvements in non-dominant hand precision drawing. Neuropsychologia, 87, 157–168. 10.1016/j.neuropsychologia.2016.05.016

Philip, B. A., Kaskutas, V., & Mackinnon, S. E. (2020). Impact of handedness on disability after unilateral upper extremity peripheral nerve disorder. HAND, 15(3), 327–334. 10.1177/1558944718810880

Philip, B. A., Li, F., Hawkins-Chernof, E., Swamidass, V., & Zwir, I. (2023). Motor Assessment With the STEGA iPad App to Measure Handwriting in Children. American Journal of Occupational Therapy, 77(3). 10.5014/ajot.2023.050098

Philip, B. A., McAvoy, M. P., & Frey, S. H. (2021). Interhemispheric Parietal-Frontal Connectivity Predicts the Ability to Acquire a Nondominant Hand Skill. Brain Connectivity, 11(4). 10.1089/brain.2020.0916

Pool, E. M., Rehme, A. K., Fink, G. R., Eickhoff, S. B., & Grefkes, C. (2014). Handedness and effective connectivity of the motor system. NeuroImage, 99, 451–460. 10.1016/j.neuroimage.2014.05.048

Potgieser, A. R., van der Hoorn, A., & de Jong, B. M. (2015). Cerebral activations related to writing and drawing with each hand. PLoS ONE, 10(5), e0126723. 10.1371/journal.pone.0126723

Power, J. D., Barnes, K. A., Snyder, A. Z., Schlaggar, B. L., & Petersen, S. E. (2012). Spurious but systematic correlations in functional connectivity MRI networks arise from subject motion. NeuroImage, 59(3), 2142–2154. 10.1016/j.neuroimage.2011.10.018

Power, J. D., Mitra, A., Laumann, T. O., Snyder, A. Z., Schlaggar, B. L., & Petersen, S. E. (2014). Methods to detect, characterize, and remove motion artifact in resting state fMRI. NeuroImage, 84, 320–341.

Reddy, N. A., Zvolanek, K. M., Moia, S., Caballero-Gaudes, C., & Bright, M. G. (2024). Denoising task-correlated head motion from motor-task fMRI data with multi-echo ICA. Imaging Neuroscience, 2. 10.1162/imag_a_00057

Saarimaki, H. (2021). Naturalistic Stimuli in Affective Neuroimaging: A Review. Front Hum Neurosci, 15, 675068. 10.3389/fnhum.2021.675068

Scarapicchia, V., Brown, C., Mayo, C., & Gawryluk, J. R. (2017). Functional Magnetic Resonance Imaging and Functional Near-Infrared Spectroscopy: Insights from Combined Recording Studies. Front Hum Neurosci, 11, 419. 10.3389/fnhum.2017.00419

Schaefer, S. Y., Haaland, K. Y., & Sainburg, R. L. (2009). Hemispheric specialization and functional impact of ipsilesional deficits in movement coordination and accuracy. Neuropsychologia, 47(13), 2953–2966. 10.1016/j.neuropsychologia.2009.06.025

Schaefer, S. Y., Mutha, P. K., Haaland, K. Y., & Sainburg, R. L. (2012). Hemispheric specialization for movement control produces dissociable differences in online corrections after stroke. Cereb Cortex, 22(6), 1407–1419. 10.1093/cercor/bhr237

Schluter, N. D., Rushworth, M. F., Passingham, R. E., & Mills, K. R. (1998). Temporary interference in human lateral premotor cortex suggests dominance for the selection of movements. A study using transcranial magnetic stimulation. Brain, 121 ( *Pt 5*), 785–799. 10.1093/brain/121.5.785

Shabbott, B. A., & Sainburg, R. L. (2008). Differentiating between two models of motor lateralization. J Neurophysiol, 100(2), 565–575. 10.1152/jn.90349.2008

Sharma, N., Pomeroy, V. M., & Baron, J.-C. (2006). Motor Imagery. Stroke, 37(7), 1941–1952. 10.1161/01.str.0000226902.43357.fc

Smith, J. C., & Frey, S. H. (2011, Feb 09). Use of independent component analysis to define regions of interest for fMRI studies. International Society for Magnetic Resonance in Medicine (ISMRM), Montreal, Canada. https://archive.ismrm.org/2011/1618.html

Smith, S. M., Jenkinson, M., Woolrich, M. W., Beckmann, C. F., Behrens, T. E. J., Johansen-Berg, H., Bannister, P. R., De Luca, M., Drobnjak, I., Flitney, D. E., Niazy, R. K., Saunders, J., Vickers, J., Zhang, Y., De Stefano, N., Brady, J. M., & Matthews, P. M. (2004). Advances in functional and structural MR image analysis and implementation as FSL. [Review]. NeuroImage, 23 *Suppl 1*, S208–219. 10.1016/j.neuroimage.2004.07.051

Stinear, C. M., Barber, P. A., Petoe, M., Anwar, S., & Byblow, W. D. (2012). The PREP algorithm predicts potential for upper limb recovery after stroke. Brain, 135(8), 2527–2535. 10.1093/brain/aws146

Stinear, C. M., & Byblow, W. D. (2004). Impaired modulation of intracortical inhibition in focal hand dystonia. Cereb Cortex, 14(5), 555–561. 10.1093/cercor/bhh017

Tam, F., Churchill, N. W., Strother, S. C., & Graham, S. J. (2011). A new tablet for writing and drawing during functional MRI. Human Brain Mapping, 32(2), 240–248.

Tsurugizawa, T., Taki, A., Zalesky, A., & Kasahara, K. (2023). Increased interhemispheric functional connectivity during non-dominant hand movement in right-handed subjects. iScience, 26(9), 107592. 10.1016/j.isci.2023.107592

Tunik, E., Ortigue, S., Adamovich, S. V., & Grafton, S. T. (2008). Differential recruitment of anterior intraparietal sulcus and superior parietal lobule during visually guided grasping revealed by electrical neuroimaging. The Journal of Neuroscience: the Official Journal of the Society for Neuroscience, 28(50), 13615–13620. 10.1523/JNEUROSCI.3303-08.2008

Tzourio-Mazoyer, N., Petit, L., Zago, L., Crivello, F., Vinuesa, N., Joliot, M., Jobard, G., Mellet, E., & Mazoyer, B. (2015). Between-hand difference in ipsilateral deactivation is associated with hand lateralization: fMRI mapping of 284 volunteers balanced for handedness. Front Hum Neurosci, 9, 5. 10.3389/fnhum.2015.00005

Valyear, K. F., & Frey, S. H. (2015). Human posterior parietal cortex mediates hand-specific planning. NeuroImage, 114, 226–238. 10.1016/j.neuroimage.2015.03.058

van Kemenade, B. M., Arikan, B. E., Podranski, K., Steinstrater, O., Kircher, T., & Straube, B. (2019). Distinct Roles for the Cerebellum, Angular Gyrus, and Middle Temporal Gyrus in Action-Feedback Monitoring. Cereb Cortex, 29(4), 1520–1531. 10.1093/cercor/bhy048

Verstynen, T., Diedrichsen, J., Albert, N., Aparicio, P., & Ivry, R. B. (2005). Ipsilateral motor cortex activity during unimanual hand movements relates to task complexity. J Neurophysiol, 93(3), 1209–1222. 10.1152/jn.00720.2004

Wang, J., Tian, Y., Wang, M., Cao, L., Wu, H., Zhang, Y., Wang, K., & Jiang, T. (2016). A lateralized top-down network for visuospatial attention and neglect. Brain Imaging Behav, 10(4), 1029–1037. 10.1007/s11682-015-9460-y

Waters-Metenier, S., Husain, M., Wiestler, T., & Diedrichsen, J. (2014). Bihemispheric Transcranial Direct Current Stimulation Enhances Effector-Independent Representations of Motor Synergy and Sequence Learning. The Journal of Neuroscience, 34(3), 1037–1050. 10.1523/jneurosci.2282-13.2014

Wischnewski, M., Kowalski, G. M., Rink, F., Belagaje, S. R., Haut, M. W., Hobbs, G., & Buetefisch, C. M. (2016). Demand on skillfulness modulates interhemispheric inhibition of motor cortices. Journal of Neurophysiology, 115(6), 2803–2813. 10.1152/jn.01076.2015

Wolpert, D. M., Goodbody, S. J., & Husain, M. (1998). Maintaining internal representations: the role of the human superior parietal lobe. Nature neuroscience, 1(6), 529–533. http://www.nature.com/neuro/journal/v1/n6/abs/nn1098_529.html

Yang, Y., Li, J., Zhao, K., Tam, F., Graham, S. J., Xu, M., & Zhou, K. (2024). Lateralized Functional Connectivity of the Sensorimotor Cortex and its Variations During Complex Visuomotor Tasks. The Journal of Neuroscience: the Official Journal of the Society for Neuroscience, 44(5). 10.1523/JNEUROSCI.0723-23.2023

Zhang, Y., Kim, J.-H., Brang, D., & Liu, Z. (2021). Naturalistic stimuli: A paradigm for multiscale functional characterization of the human brain. Current Opinion in Biomedical Engineering, 19, 100298. 10.1016/j.cobme.2021.100298

