## Supplementary material for "Neural mechanisms of handedness for precision drawing: hand-dependent engagement of cortical networks for bimanual control": SuppFigure1_BOLD_distrib.pdf

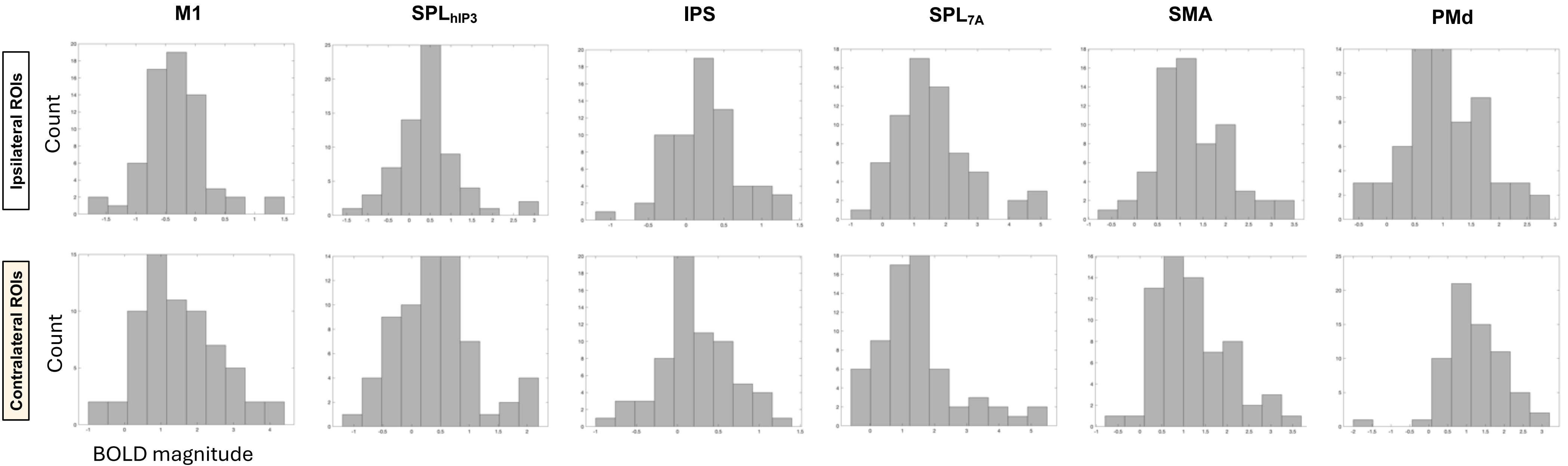


**Supplementary Figure 1**: Histograms of BOLD magnitudes (Draw > Rest) across participants for each ROI, showing normal or approximately normal distributions.
