## Supplementary material for "Neural mechanisms of handedness for precision drawing: hand-dependent engagement of cortical networks for bimanual control": Supplementary_Text_clean.docx

**S1. Methods**

**S1.1. fMRIPrep Preprocessing**

**S1.1.1. Preprocessing of B0 inhomogeneity mappings:** A B0-nonuniformity map (or fieldmap) was estimated based on two echo-planar imaging (EPI) references with topup (Andersson et al., 2003).

**S1.1.2. Anatomical preprocessing:** T1-weighted (T1w) image was corrected for intensity non-uniformity (INU) with N4BiasFieldCorrection (Tustison et al., 2010), distributed with ANTs 2.5.3 (Avants et al., 2008; RRID:SCR_004757), and used as T1w-reference throughout the workflow. The T1w-reference was then skull-stripped with a Nipype implementation of the antsBrainExtraction.sh workflow (from ANTs), using OASIS30ANTs as target template. Brain tissue segmentation of cerebrospinal fluid (CSF), white-matter (WM) and gray-matter (GM) was performed on the brain-extracted T1w using fast (FSL, Zhang et al., 2002; RRID:SCR_002823). Brain surfaces were reconstructed using recon-all ((FreeSurfer 7.3.2, Dale et al., 1999; RRID:SCR_001847), and the brain mask estimated previously was refined with a custom variation of the method to reconcile ANTs-derived and FreeSurfer-derived segmentations of the cortical gray-matter of Mindboggle (Klein et al., 2017; RRID:SCR_002438). A T2-weighted image was used to improve pial surface refinement. Brain surfaces were reconstructed using recon-all (FreeSurfer 7.3.2), and the brain mask estimated previously was refined with a custom variation of the method to reconcile ANTs-derived and FreeSurfer-derived segmentations of the cortical gray-matter of Mindboggle. Volume-based spatial normalization was performed to output space MNI ICBM 152 Nonlinear Symmetrical template version 2009c (Fonov et al., 2009; RRIC:SCR_008796; TemplateFlow ID: MNI152NLin2009cSym) at 2mm3 resolution, through nonlinear registration with antsRegistration (ANTs 2.5.3), using brain-extracted versions of both T1w reference and the T1w template.

**S1.1.3. Functional preprocessing:** For each of the 6 BOLD runs per subject, the following preprocessing was performed. First, a reference volume was generated, using a custom methodology of fMRIPrep, for use in head motion correction. Head-motion parameters with respect to the BOLD reference (transformation matrices, and six corresponding rotation and translation parameters) are estimated before any spatiotemporal filtering using mcflirt (FSL, Jenkinson et al., 2002). The estimated fieldmap was then aligned with rigid-registration to the target EPI (echo-planar imaging) reference run. The field coefficients were mapped on to the reference EPI using the transform. The BOLD reference was then co-registered to the T1w reference using bbregister (FreeSurfer) which implements boundary-based registration (Freesurfer, Greve & Fischl, 2009). Co-registration was configured with six degrees of freedom. The aligned T2w image was used for initial co-registration. Several confounding time-series were calculated based on the preprocessed BOLD: framewise displacement (FD), and global signal extracted from the white matter (WM), cerebrospinal fluid (CSF), and whole-brain masks. FD was computed using fsl_motion_outliers (relative root mean square displacement between affines, Jenkinson et al., 2002). Additionally, a set of physiological regressors were extracted to allow for component-based noise correction (CompCor, Behzadi et al., 2007). Principal components were estimated after high-pass filtering the preprocessed BOLD time-series (using a discrete cosine filter with 128s cut-off) for the two CompCor variants: temporal (tCompCor) and anatomical (aCompCor). tCompCor components were then calculated from the top 2% variable voxels within the brain mask. For aCompCor, three probabilistic masks (CSF, WM and combined CSF+WM) were generated in anatomical space. The implementation differs from that of Behzadi et al. in that instead of eroding the masks by 2 pixels on BOLD space, a mask of pixels that likely contain a volume fraction of GM was subtracted from the aCompCor masks. This mask was obtained by dilating a GM mask extracted from the FreeSurfer’s aseg segmentation, and it ensures components are not extracted from voxels containing a minimal fraction of GM. Finally, these masks were resampled into BOLD space and binarized by thresholding at 0.99 (as in the original implementation). Components were also calculated separately within the WM and CSF masks. For each CompCor decomposition, the k components with the largest singular values were retained, such that the retained components’ time series were sufficient to explain 50 percent of variance across the nuisance mask (CSF, WM, combined, or temporal). The remaining components were dropped from consideration. The head-motion estimates calculated in the correction step were also included as confounds. The confound time series derived from head motion estimates and global signals were expanded with the inclusion of temporal derivatives and quadratic terms for each (Satterthwaite et al., 2013). Additional nuisance timeseries were calculated by means of principal components analysis of the signal found within a thin band (crown) of voxels around the edge of the brain, as proposed by Patriat et al. (2017). All resamplings were performed with a single interpolation step by composing all the pertinent transformations (i.e. head-motion transform matrices, susceptibility distortion correction, and co-registrations to anatomical and output spaces). Gridded (volumetric) resamplings were performed using nitransforms, configured with cubic B-spline interpolation.

**S.1.4. fMRI Motion Outlier Identification and Screening**

**S.1.4.1. Screening for High Motion**. To screen participants with high motion, two screening criteria were employed: 1) Participant-level FD threshold: participant-mean FD was calculated by averaging FD within each run, and then averaging across runs for each participant. Participants were removed from analysis if their participant-mean FD was > 0.5 mm (Power et al., 2012; Power et al., 2014); based on this criterion, 1 participant was excluded. 2) Within-participant motion outliers: for each participant, outlier volumes were defined as FD > 75^th^ percentile + (1.5 * inter-quartile range). This threshold identified 4.6% of volumes as outliers (# outlier volumes per scan: range 0-68, mean 22.4, quartiles 10, 20, 33). A participant would be excluded if their # outliers exceeded 20% of the data in any scan (98 volumes), but no participants met that criterion.

**S.1.4.1. Screening for Excess Task-Correlated Motion**. To screen participants with excess task-correlated motion, the coefficient of determination (Pearson r^2^) was calculated between FD and the task timecourse. Correlation was calculated for each run, and averaged across runs within each participant. These values ranged 0.00004-0.075 (mean 0.021, median 0.016). This process was repeated for the first derivative of the task timecourse, leading to correlation values that ranged 0.001-0.137 (mean 0.047, median 0.033). Participants were excluded from analysis if their correlation value exceeded the outlier threshold of 75^th^ percentile + (1.5 * inter-quartile range) for either the timecourse (threshold r^2^ = 0.161) or its derivative (threshold r^2^ = 0.239). Based on this criterion, 2 participants were excluded.

**S.1.4.3. Screening for Task Non-Compliance**. To screen participants who failed to comply with the task, a two-step process was used. First, experimenters qualitatively evaluated participant compliance during data collection (a yes/no warning flag); 7 participants were flagged for follow-up. Second, the flagged participants were explored using objective data for 3 measures of task compliance: number of times they simultaneously touched the tablet with the stylus and a finger (which cause the drawn line to jump across the screen), amount of time spent drawing during drawing blocks (which would be low if the participant was slow to respond to the change from rest to draw), and performance (BIS, see *Supplementary text S1.6. Performance Data Processing*). One flagged participant had the highest number of touches of all participants, and also had the 3^rd^-lowest performance, so this participant was excluded from analysis. No other participants showed atypical task compliance, though 3/7 flagged participants had been excluded due to high participant-level FD or task-correlated motion as described above. The remaining 3/7 flagged participants did not show atypical task compliance by any objective metric and thus were retained in the sample.

**S1.5. Performance Data Collection.** Pen position was recorded at 60 Hz during drawing blocks of the fMRI task. To remove artifacts caused by the participant accidentally touching the tablet with a finger (which caused the recorded location to “jump” instantly across the tablet), raw data points were removed if their velocity exceeded the 4^th^ standard deviation, calculated within each scan.

**S1.6. Performance Data Processing:** Based on the position data, six performance measures were calculated: position accuracy (“posErr” for brevity; -1 * position error, mm), smoothness “velSm” (“velSm;” -1 * number of velocity peaks per 45 mm shape part), direction accuracy “dirAcc” (“dirAcc;” -1 * direction error, degrees), velocity (“spd” to avoid confusion with velSm; root mean squared, mm / s), and two secondary measures: Balanced Integrated Score (BIS, “bis”) and Speed Accuracy Tradeoff (SAT, “sat”), detailed in the following paragraphs.

BIS captures overall performance by integrating speed and accuracy measures (Liesefeld & Janczyk, 2023; Vandierendonck, 2021). BIS is calculated for each participant-hand pair (i.e. each participant had a LH and a RH value) as zscore(speed) + zscore(position accuracy), using participant-hand averages for speed and error, and z-scoring across the entire dataset (all participants & hands). An alternative calculation of BIS was explored using direction accuracy rather than position accuracy, but the two measures were highly correlated (r^2^ > 0.9), so only position accuracy was used.

SAT captures participant tradeoffs between fast (feedforward) and slow corrective (feedback) actions, to assess the participant’s relative reliance on the two lateralized components in the complementary dominance hypothesis: the left-hemisphere feedforward specialization and right-hemisphere feedback specialization (Sainburg, 2002; Wang & Sainburg, 2007). Following our calculation of BIS, we determined SAT as zscore(speed) – zscore(position accuracy), such that a high SAT value will reflect a fast imprecise movement.

Performance variables were initially generated at the participant-hand level, but for each participant the LH and RH values were highly colinear, reflecting the existence of a “bilateral/overall dexterity” participant trait (**Table S1 column 1**). To enable analysis of left/right performance differences without collinearity, an “asymmetry” value (“Asym”) was calculated for each participant and variable as (RH-LH)/(RH+LH) (after offset to minimum = 0), such that high values represent the advantage of the preferred RH (Vingerhoets et al., 2023). Asymmetry’s correlations with LH and RH were uniformly lower than the correlation between LH and RH, as shown in **Table S1** **columns 2-3**.

| **Table S1:** Correlations between performance variables across hands. | | | |
| --- | --- | --- | --- |
|  | **Coefficient of determination (r^2^)** | | |
| **Measure** | **LH : RH** | **Asym : LH** | **Asym : RH** |
| Position Accuracy | 0.357 | 0.038 | 0.427 |
| Velocity Smoothness | 0.829 | 0.009 | 0.228 |
| Direction Accuracy | 0.564 | 0.101 | 0.148 |
| Velocity | 0.856 | 0.003 | 0.101 |
| SAT | 0.865 | 0.000 | 0.112 |
| BIS | 0.826 | 0.036 | 0.048 |

The “bilateral/overall dexterity” trait also manifested as across-measure correlations, which were addressed by minimizing the number of measures (*S1.4* below) or using a stepwise process to avoid collinearities (*S1.5* below).

**S1.7. Performance Confound Analysis (Exploratory GLME #1):** First, to assess the effect of performance confounds on BOLD magnitude in each ROI (i.e. main manuscript section 2.7.4, *magnitude ROI analysis*), the main generalized linear mixed effects (GLME) analysis was repeated with the addition of 2 performance measurements. First, to control for amount of movement, velocity (“vel;” root mean squared mm / s) was included. Second, to control for difficulty, velocity smoothness (“smooth;” -1 * number of velocity peaks per 45 mm shape part) was included. Velocity smoothness was chosen because a previous study identified it as the most hand-specific performance variable in a similar precision drawing task (Philip & Frey, 2016).

In this performance confound analysis, each measurement was coded as Mean and Asym variables, leading to 4 candidate performance variables.

To limit the number of factors in each model, a screening procedure was used to select 2 factors from the 4 candidates, separately for each ROI. The screening procedure entailed: (1) for each candidate variable, modeling ROI magnitude with two univariable GLMs, one for hand_RH_ and one for hand_LH_; (2) defining the candidate variable’s strength from its GLM T-statistics as $strength={mean}_{hands}\left| T \right|$; (3) ranking the candidate variables by strength; and (4) selecting the strongest 2 candidate variables as factors in the main GLME.

Model evaluation and statistical tests were performed as described in the main analysis. False discovery rate (FDR) correction was performed separately once for the primary hypothesis (hand_LH-RH_; 12 values, one per GLME), secondary hypothesis (group and hand_LH-RH_*group; 24 values, 2 per GLME), participant characteristics (24 values, 2 per GLME), and performance factors (24 values, 2 per GLME).

**S1.8. Performance Influence Analysis (Exploratory GLME #2): To** explore possible behavior factors that might influence BOLD magnitude in each ROI, an additional GLME analysis was performed using the repeated-measures factor Subject, hand_LH-RH_, and factors drawn from 21 candidates: age, sex, group, and 18 performance variables. The 18 performance variables reflected 6 measurements (velocity, velocity smoothness, position accuracy, direction accuracy, BIS, SAT) * 3 values (Asym, LH, RH). This differred from the previous analysis because LH and RH were included to identify hand-specific factors, which required the omission of “Mean” because of the high correlations between LH:Mean and RH:Mean (r > 0.9).

For each ROI, the model factors were selected via a 3-step screening procedure. Steps (1) and (2) were identical to *S1.4* above, but then (3) a stepwise procedure was used to maximize the number of factors while avoiding collinearity. In this stepwise procedure, candidates were tested in order of strength, and the candidate was added to the model if it was not correlated with any factors already in the model (Pearson’s *r*, α = 0.05/# of correlations tested).

Model evaluation and statistical tests were performed as described in the main analysis. FDR False discovery rate correction was performed as described above (*S1.4*), except the quantity of participant characteristics and performance factors was determined by the stepwise process in the previous paragraph.

**S2. Results**

**S2.1. Performance Confound Analysis:** In our analysis of performance confounds on our main ROI analysis, no performance factor contributed significantly to BOLD magnitude (p-uncorrected > 0.11), and the inclusion of performance confounds did not change the results of hypothesized effects (hand_LH-RH_, group, hand_LH-RH_ * group), as shown in **Table S2**.

| **Table S2**. Performance confound analysis, GLME factors. | | | | |
| --- | --- | --- | --- | --- |
| **ROI** | **Factor** | **T** | **p-uncorrected** | **p-FDR** |
| M1_Ipsi | hand(LH-RH) | 2.698 | 0.009 | 0.022* |
| M1_Ipsi | isPatient | -0.148 | 0.883 | 0.955 |
| M1_Ipsi | sexF | 0.966 | 0.338 | 0.941 |
| M1_Ipsi | DASH_ability | 0.085 | 0.932 | 0.959 |
| M1_Ipsi | age | 0.484 | 0.630 | 0.941 |
| M1_Ipsi | velSm_Mean | -0.571 | 0.570 | 0.940 |
| M1_Ipsi | spd_Mean | 0.367 | 0.715 | 0.940 |
| M1_Ipsi | hand(LH-RH) * isPatient | -0.649 | 0.519 | 0.922 |
| M1_Contra | hand(LH-RH) | -3.807 | 0.000 | 0.002* |
| M1_Contra | isPatient | -1.282 | 0.205 | 0.922 |
| M1_Contra | age | -1.673 | 0.100 | 0.855 |
| M1_Contra | sexF | -0.188 | 0.851 | 0.941 |
| M1_Contra | DASH_ability | -1.340 | 0.185 | 0.941 |
| M1_Contra | velSm_Asym | -0.421 | 0.675 | 0.940 |
| M1_Contra | velSm_Mean | 1.301 | 0.198 | 0.680 |
| M1_Contra | hand(LH-RH) * isPatient | 0.673 | 0.504 | 0.922 |
| SPL_hIP3__Ipsi | hand(LH-RH) | -4.010 | 0.000 | 0.002* |
| SPL _hIP3__Ipsi | isPatient | 0.208 | 0.836 | 0.955 |
| SPL _hIP3__Ipsi | sexF | 1.151 | 0.254 | 0.941 |
| SPL _hIP3__Ipsi | DASH_ability | 0.145 | 0.886 | 0.941 |
| SPL _hIP3__Ipsi | age | 0.516 | 0.608 | 0.941 |
| SPL _hIP3__Ipsi | velSm_Mean | -1.569 | 0.122 | 0.587 |
| SPL _hIP3__Ipsi | spd_Mean | 0.373 | 0.710 | 0.940 |
| SPL _hIP3__Ipsi | hand(LH-RH) * isPatient | 0.944 | 0.349 | 0.922 |
| SPL _hIP3__Contra | hand(LH-RH) | 1.521 | 0.134 | 0.224 |
| SPL _hIP3__Contra | isPatient | 0.817 | 0.417 | 0.922 |
| SPL _hIP3__Contra | sexF | 1.245 | 0.218 | 0.941 |
| SPL _hIP3__Contra | age | 0.526 | 0.601 | 0.941 |
| SPL _hIP3__Contra | DASH_ability | 0.759 | 0.451 | 0.941 |
| SPL _hIP3__Contra | velSm_Mean | -1.140 | 0.259 | 0.777 |
| SPL _hIP3__Contra | velSm_Asym | 0.322 | 0.749 | 0.940 |
| SPL _hIP3__Contra | hand(LH-RH) * isPatient | 0.057 | 0.955 | 0.955 |
| IPS_Ipsi | hand(LH-RH) | -3.026 | 0.004 | 0.011* |
| IPS_Ipsi | isPatient | -0.716 | 0.477 | 0.922 |
| IPS_Ipsi | sexF | 2.037 | 0.046 | 0.855 |
| IPS_Ipsi | DASH_ability | -1.514 | 0.135 | 0.855 |
| IPS_Ipsi | age | 0.587 | 0.560 | 0.941 |
| IPS_Ipsi | velSm_Mean | -0.891 | 0.377 | 0.904 |
| IPS_Ipsi | spd_Mean | 1.619 | 0.111 | 0.587 |
| IPS_Ipsi | hand(LH-RH) * isPatient | -0.492 | 0.625 | 0.922 |
| IPS_Contra | hand(LH-RH) | 1.462 | 0.149 | 0.224 |
| IPS_Contra | isPatient | -1.219 | 0.228 | 0.922 |
| IPS_Contra | sexF | 1.756 | 0.085 | 0.855 |
| IPS_Contra | DASH_ability | -1.539 | 0.129 | 0.855 |
| IPS_Contra | age | 0.594 | 0.555 | 0.941 |
| IPS_Contra | velSm_Mean | -0.942 | 0.350 | 0.904 |
| IPS_Contra | spd_Mean | 1.324 | 0.191 | 0.680 |
| IPS_Contra | hand(LH-RH) * isPatient | 1.011 | 0.316 | 0.922 |
| SPL_7A__Ipsi | hand(LH-RH) | -1.251 | 0.216 | 0.288 |
| SPL_7A__Ipsi | isPatient | 0.355 | 0.724 | 0.922 |
| SPL_7A__Ipsi | age | 0.285 | 0.777 | 0.941 |
| SPL_7A__Ipsi | sexF | 0.387 | 0.700 | 0.941 |
| SPL_7A__Ipsi | DASH_ability | -0.001 | 0.999 | 0.999 |
| SPL_7A__Ipsi | velSm_Asym | 1.586 | 0.118 | 0.587 |
| SPL_7A__Ipsi | velSm_Mean | -0.141 | 0.888 | 0.940 |
| SPL_7A__Ipsi | hand(LH-RH) * isPatient | 0.401 | 0.690 | 0.922 |
| SPL_7A__Contra | hand(LH-RH) | -0.979 | 0.332 | 0.351 |
| SPL_7A__Contra | isPatient | -0.546 | 0.587 | 0.922 |
| SPL_7A__Contra | sexF | 1.487 | 0.142 | 0.855 |
| SPL_7A__Contra | DASH_ability | -0.514 | 0.610 | 0.941 |
| SPL_7A__Contra | age | 0.587 | 0.559 | 0.941 |
| SPL_7A__Contra | velSm_Mean | -0.359 | 0.721 | 0.940 |
| SPL_7A__Contra | spd_Mean | 1.572 | 0.122 | 0.587 |
| SPL_7A__Contra | hand(LH-RH) * isPatient | 0.964 | 0.339 | 0.922 |
| PMd_Ipsi | hand(LH-RH) | -0.948 | 0.347 | 0.351 |
| PMd_Ipsi | isPatient | -0.495 | 0.622 | 0.922 |
| PMd_Ipsi | sexF | 0.729 | 0.469 | 0.941 |
| PMd_Ipsi | age | -0.720 | 0.474 | 0.941 |
| PMd_Ipsi | DASH_ability | -0.234 | 0.816 | 0.941 |
| PMd_Ipsi | velSm_Mean | -0.208 | 0.836 | 0.940 |
| PMd_Ipsi | velSm_Asym | -0.231 | 0.818 | 0.940 |
| PMd_Ipsi | hand(LH-RH) * isPatient | 1.502 | 0.139 | 0.922 |
| PMd_Contra | hand(LH-RH) | -0.940 | 0.351 | 0.351 |
| PMd_Contra | isPatient | 0.074 | 0.941 | 0.955 |
| PMd_Contra | sexF | 0.710 | 0.480 | 0.941 |
| PMd_Contra | age | -0.570 | 0.571 | 0.941 |
| PMd_Contra | DASH_ability | -0.141 | 0.888 | 0.941 |
| PMd_Contra | velSm_Mean | -0.262 | 0.794 | 0.940 |
| PMd_Contra | velSm_Asym | -0.064 | 0.949 | 0.949 |
| PMd_Contra | hand(LH-RH) * isPatient | -0.347 | 0.730 | 0.922 |
| SMA_Ipsi | hand(LH-RH) | 3.529 | 0.001 | 0.003* |
| SMA_Ipsi | isPatient | -0.747 | 0.458 | 0.922 |
| SMA_Ipsi | DASH_ability | -0.222 | 0.825 | 0.941 |
| SMA_Ipsi | age | 0.423 | 0.674 | 0.941 |
| SMA_Ipsi | sexF | 0.372 | 0.711 | 0.941 |
| SMA_Ipsi | spd_Mean | 0.227 | 0.822 | 0.940 |
| SMA_Ipsi | velSm_Asym | -0.125 | 0.901 | 0.940 |
| SMA_Ipsi | hand(LH-RH) * isPatient | 1.367 | 0.177 | 0.922 |
| SMA_Contra | hand(LH-RH) | -1.623 | 0.110 | 0.220 |
| SMA_Contra | isPatient | -1.101 | 0.276 | 0.922 |
| SMA_Contra | age | 0.288 | 0.774 | 0.941 |
| SMA_Contra | sexF | 0.249 | 0.804 | 0.941 |
| SMA_Contra | DASH_ability | -0.947 | 0.348 | 0.941 |
| SMA_Contra | velSm_Asym | 0.390 | 0.698 | 0.940 |
| SMA_Contra | velSm_Mean | -1.711 | 0.092 | 0.587 |
| SMA_Contra | hand(LH-RH) * isPatient | -0.114 | 0.910 | 0.955 |
| *Note*: * = p-FDR < 0.05. | | | | |

**S2.2. Performance Influence Analysis:** In our broad analysis of possible performance effects, no factor other than hand_LH-RH_ contributed significantly to BOLD magnitude, as shown in **Table S3**. We found a non-significant trends toward effects of RH position accuracy in M1-contra (p-FDR = 0.055) and IPS-ipsi (p-FDR = 0.086); all other factors p-FDR > 0.17.

| **Table S3**. Performance influence analysis, GLME factors. | | | | |
| --- | --- | --- | --- | --- |
| **ROI** | **Factor** | **T** | **p-uncorrected** | **p-FDR** |
| M1_Ipsi | hand(LH-RH) | 2.786 | 0.007 | 0.017* |
| M1_Ipsi | SAT_Asym | -2.244 | 0.029 | 0.303 |
| M1_Ipsi | dirAcc_RH | -1.607 | 0.113 | 0.501 |
| M1_Ipsi | sexF | 1.302 | 0.198 | 0.755 |
| M1_Ipsi | isPatient | -1.555 | 0.125 | 0.415 |
| M1_Ipsi | SAT_RH | 0.533 | 0.596 | 0.873 |
| M1_Contra | hand(LH-RH) | -4.088 | 0.000 | 0.001* |
| M1_Contra | posAcc_RH | 3.455 | 0.001 | 0.055 |
| M1_Contra | SAT_Asym | -2.120 | 0.038 | 0.312 |
| M1_Contra | velSm_LH | 2.665 | 0.010 | 0.176 |
| M1_Contra | BIS_LH | 0.399 | 0.691 | 0.916 |
| M1_Contra | dirAcc_Asym | -0.029 | 0.977 | 0.986 |
| M1_Contra | isPatient | -0.342 | 0.733 | 0.733 |
| SPL _hIP3__Ipsi | hand(LH-RH) | -4.125 | 0.000 | 0.001* |
| SPL _hIP3__Ipsi | velSm_RH | -1.461 | 0.149 | 0.528 |
| SPL _hIP3__Ipsi | posAcc_Asym | 1.184 | 0.241 | 0.650 |
| SPL _hIP3__Ipsi | sexF | 0.715 | 0.478 | 0.796 |
| SPL _hIP3__Ipsi | DASH_ability | -0.248 | 0.805 | 0.905 |
| SPL _hIP3__Ipsi | SAT_LH | 0.094 | 0.926 | 0.986 |
| SPL _hIP3__Ipsi | dirAcc_LH | -0.057 | 0.955 | 0.986 |
| SPL _hIP3__Contra | hand(LH-RH) | 1.859 | 0.068 | 0.102 |
| SPL _hIP3__Contra | posAcc_Asym | 1.214 | 0.230 | 0.650 |
| SPL _hIP3__Contra | velSm_RH | -0.944 | 0.349 | 0.658 |
| SPL _hIP3__Contra | sexF | 0.868 | 0.389 | 0.796 |
| SPL _hIP3__Contra | dirAcc_LH | -0.544 | 0.588 | 0.873 |
| SPL _hIP3__Contra | spd_LH | -0.065 | 0.948 | 0.986 |
| SPL _hIP3__Contra | DASH_ability | 0.390 | 0.698 | 0.905 |
| IPS_Ipsi | hand(LH-RH) | -3.935 | 0.000 | 0.001* |
| IPS_Ipsi | posAcc_RH | 3.071 | 0.003 | 0.086 |
| IPS_Ipsi | DASH_ability | -1.409 | 0.164 | 0.755 |
| IPS_Ipsi | SAT_Asym | -0.891 | 0.377 | 0.666 |
| IPS_Ipsi | BIS_LH | 0.990 | 0.326 | 0.658 |
| IPS_Ipsi | dirAcc_Asym | 0.223 | 0.824 | 0.986 |
| IPS_Ipsi | velSm_LH | 0.456 | 0.650 | 0.907 |
| IPS_Contra | hand(LH-RH) | 2.381 | 0.021 | 0.041* |
| IPS_Contra | posAcc_RH | 2.490 | 0.016 | 0.207 |
| IPS_Contra | DASH_ability | -1.424 | 0.160 | 0.755 |
| IPS_Contra | SAT_Asym | -0.969 | 0.336 | 0.658 |
| IPS_Contra | BIS_LH | 0.713 | 0.478 | 0.754 |
| IPS_Contra | velSm_LH | 0.264 | 0.793 | 0.986 |
| IPS_Contra | dirAcc_Asym | -0.081 | 0.935 | 0.986 |
| SPL_7A__Ipsi | hand(LH-RH) | -1.231 | 0.223 | 0.268 |
| SPL_7A__Ipsi | posAcc_RH | 1.649 | 0.105 | 0.501 |
| SPL_7A__Ipsi | BIS_RH | 1.487 | 0.142 | 0.528 |
| SPL_7A__Ipsi | dirAcc_Asym | 0.969 | 0.337 | 0.658 |
| SPL_7A__Ipsi | DASH_ability | -0.756 | 0.452 | 0.796 |
| SPL_7A__Ipsi | velSm_LH | -0.175 | 0.862 | 0.986 |
| SPL_7A__Ipsi | SAT_Asym | 0.193 | 0.848 | 0.986 |
| SPL_7A__Contra | hand(LH-RH) | -0.530 | 0.598 | 0.652 |
| SPL_7A__Contra | posAcc_RH | 2.023 | 0.048 | 0.316 |
| SPL_7A__Contra | BIS_RH | 1.302 | 0.198 | 0.617 |
| SPL_7A__Contra | dirAcc_Asym | 0.514 | 0.609 | 0.873 |
| SPL_7A__Contra | DASH_ability | -0.905 | 0.369 | 0.796 |
| SPL_7A__Contra | SAT_Asym | 0.320 | 0.750 | 0.969 |
| SPL_7A__Contra | velSm_LH | 0.070 | 0.945 | 0.986 |
| PMd_Ipsi | hand(LH-RH) | -0.140 | 0.889 | 0.889 |
| PMd_Ipsi | posAcc_Asym | 2.088 | 0.041 | 0.312 |
| PMd_Ipsi | sexF | 1.255 | 0.214 | 0.755 |
| PMd_Ipsi | BIS_LH | 1.114 | 0.270 | 0.650 |
| PMd_Ipsi | BIS_Asym | -0.922 | 0.360 | 0.658 |
| PMd_Ipsi | DASH_ability | 0.303 | 0.763 | 0.905 |
| PMd_Ipsi | velSm_LH | 0.081 | 0.936 | 0.986 |
| PMd_Contra | hand(LH-RH) | -1.353 | 0.181 | 0.242 |
| PMd_Contra | posAcc_Asym | 1.939 | 0.057 | 0.338 |
| PMd_Contra | sexF | 1.158 | 0.252 | 0.755 |
| PMd_Contra | BIS_LH | 1.127 | 0.264 | 0.650 |
| PMd_Contra | BIS_Asym | -0.819 | 0.416 | 0.702 |
| PMd_Contra | DASH_ability | 0.197 | 0.844 | 0.905 |
| PMd_Contra | velSm_LH | -0.018 | 0.986 | 0.986 |
| SMA_Ipsi | hand(LH-RH) | 4.990 | 0.000 | 0.000* |
| SMA_Ipsi | posAcc_Asym | 1.343 | 0.185 | 0.612 |
| SMA_Ipsi | velSm_LH | -1.541 | 0.129 | 0.525 |
| SMA_Ipsi | spd_Asym | -0.935 | 0.354 | 0.658 |
| SMA_Ipsi | isPatient | -1.022 | 0.311 | 0.415 |
| SMA_Ipsi | sexF | 0.095 | 0.924 | 0.924 |
| SMA_Ipsi | BIS_LH | 0.705 | 0.484 | 0.754 |
| SMA_Contra | hand(LH-RH) | -2.019 | 0.048 | 0.083 |
| SMA_Contra | posAcc_Asym | 1.761 | 0.084 | 0.443 |
| SMA_Contra | velSm_LH | -1.163 | 0.250 | 0.650 |
| SMA_Contra | spd_Asym | -0.805 | 0.424 | 0.702 |
| SMA_Contra | isPatient | -1.262 | 0.212 | 0.415 |
| SMA_Contra | sexF | 0.511 | 0.612 | 0.905 |
| SMA_Contra | BIS_LH | 0.943 | 0.350 | 0.658 |
| SMA_Contra | dirAcc_RH | -0.418 | 0.678 | 0.916 |
| *Note*: * = p-FDR < 0.05. | | | | |

**References for Supplementary Text**

Andersson, J. L., Skare, S., & Ashburner, J. (2003). How to correct susceptibility distortions in spin-echo echo-planar images: application to diffusion tensor imaging. *NeuroImage*, *20*(2), 870-888. <https://doi.org/10.1016/S1053-8119(03)00336-7>

Avants, B. B., Epstein, C. L., Grossman, M., & Gee, J. C. (2008). Symmetric diffeomorphic image registration with cross-correlation: evaluating automated labeling of elderly and neurodegenerative brain. *Medical image analysis*, *12*(1), 26-41.

Behzadi, Y., Restom, K., Liau, J., & Liu, T. T. (2007). A component based noise correction method (CompCor) for BOLD and perfusion based fMRI. *NeuroImage*, *37*(1), 90-101. <https://doi.org/10.1016/j.neuroimage.2007.04.042>

Dale, A. M., Fischl, B., & Sereno, M. I. (1999). Cortical surface-based analysis: I. Segmentation and surface reconstruction. *NeuroImage*, *9*(2), 179-194.

Fonov, V. S., Evans, A. C., McKinstry, R. C., Almli, C. R., & Collins, D. (2009). Unbiased nonlinear average age-appropriate brain templates from birth to adulthood. *NeuroImage*(47), S102.

Greve, D. N., & Fischl, B. (2009). Accurate and robust brain image alignment using boundary-based registration. [Evaluation Study]. *NeuroImage*, *48*(1), 63-72. <https://doi.org/10.1016/j.neuroimage.2009.06.060>

Jenkinson, M., Bannister, P., Brady, M., & Smith, S. (2002). Improved optimization for the robust and accurate linear registration and motion correction of brain images. *NeuroImage*, *17*(2), 825-841.

Klein, A., Ghosh, S. S., Bao, F. S., Giard, J., Häme, Y., Stavsky, E., Lee, N., Rossa, B., Reuter, M., & Chaibub Neto, E. (2017). Mindboggling morphometry of human brains. *PLoS Computational Biology*, *13*(2), e1005350.

Liesefeld, H. R., & Janczyk, M. (2023). Same same but different: Subtle but consequential differences between two measures to linearly integrate speed and accuracy (LISAS vs. BIS). *Behav Res Methods*, *55*(3), 1175-1192. <https://doi.org/10.3758/s13428-022-01843-2>

Patriat, R., Reynolds, R. C., & Birn, R. M. (2017). An improved model of motion-related signal changes in fMRI. *NeuroImage*, *144*, 74-82.

Philip, B. A., & Frey, S. H. (2016). Increased functional connectivity between cortical hand areas and praxis network associated with training-related improvements in non-dominant hand precision drawing. *Neuropsychologia*, *87*, 157-168. <https://doi.org/10.1016/j.neuropsychologia.2016.05.016>

Power, J. D., Barnes, K. A., Snyder, A. Z., Schlaggar, B. L., & Petersen, S. E. (2012). Spurious but systematic correlations in functional connectivity MRI networks arise from subject motion. *NeuroImage*, *59*(3), 2142-2154. <https://doi.org/10.1016/j.neuroimage.2011.10.018>

Power, J. D., Mitra, A., Laumann, T. O., Snyder, A. Z., Schlaggar, B. L., & Petersen, S. E. (2014). Methods to detect, characterize, and remove motion artifact in resting state fMRI. *NeuroImage*, *84*, 320-341.

Sainburg, R. L. (2002). Evidence for a dynamic-dominance hypothesis of handedness. *Experimental brain research Experimentelle Hirnforschung Expérimentation cérébrale*, *142*(2), 241-258. <https://doi.org/10.1007/s00221-001-0913-8>

Satterthwaite, T. D., Elliott, M. A., Gerraty, R. T., Ruparel, K., Loughead, J., Calkins, M. E., Eickhoff, S. B., Hakonarson, H., Gur, R. C., & Gur, R. E. (2013). An improved framework for confound regression and filtering for control of motion artifact in the preprocessing of resting-state functional connectivity data. *NeuroImage*, *64*, 240-256.

Tustison, N. J., Avants, B. B., Cook, P. A., Zheng, Y., Egan, A., Yushkevich, P. A., & Gee, J. C. (2010). N4ITK: improved N3 bias correction. *IEEE transactions on medical imaging*, *29*(6), 1310-1320.

Vandierendonck, A. (2021). On the Utility of Integrated Speed-Accuracy Measures when Speed-Accuracy Trade-off is Present. *J Cogn*, *4*(1), 22. <https://doi.org/10.5334/joc.154>

Vingerhoets, G., Verhelst, H., Gerrits, R., Badcock, N., Bishop, D. V., Carey, D., Flindall, J., Grimshaw, G., Harris, L. J., & Hausmann, M. (2023). Laterality indices consensus initiative (LICI): A Delphi expert survey report on recommendations to record, assess, and report asymmetry in human behavioural and brain research. *Laterality*, *28*(2-3), 122-191.

Wang, J., & Sainburg, R. L. (2007). The dominant and nondominant arms are specialized for stabilizing different features of task performance. *Experimental brain research Experimentelle Hirnforschung Expérimentation cérébrale*, *178*(4), 565-570. <https://doi.org/10.1007/s00221-007-0936-x>

Zhang, Y., Brady, M., & Smith, S. (2002). Segmentation of brain MR images through a hidden Markov random field model and the expectation-maximization algorithm. *IEEE transactions on medical imaging*, *20*(1), 45-57.
